# Cholesterol efflux protein, ABCA1, supports anti-cancer functions of myeloid immune cells

**DOI:** 10.1101/2025.02.19.638515

**Authors:** Shruti V. Bendre, Yu Wang, Basel Hajyousif, K C Rajendra, Shounak G. Bhogale, Dhanya Pradeep, Natalia Krawczynska, Claire P. Schane, Erin Weisser, Avni Singh, Simon Han, Hannah Kim, Lara Kockaya, Anasuya Das Gupta, Adam T. Nelczyk, Hashni Epa Vidana Gamage, Yifan Fei, Xingyu Guo, Ryan J. Deaton, Maria Sverdlov, Peter H. Gann, Saurabh Sinha, Kun Wang, Kevin Van Bortle, Emad Tajkorshid, Wendy A. Woodward, Wonhwa Cho, Erik R. Nelson

## Abstract

Although immune therapy has seen significant advances, the majority of breast and other solid tumors do not respond or quickly develop *de novo* resistance. One factor driving resistance is highly immune suppressive myeloid cells (MCs) such as macrophages. Previous work has established clinical links between cholesterol and cancer outcome, and that MC function can be regulated through disruption in cholesterol metabolism. Thus, we screened for proteins that were expressed in MCs, involved in cholesterol homeostasis and whose expression was associated with survival; we identify the cholesterol efflux protein ABCA1. Preclinical studies revealed that ABCA1 activity resulted in increased anti-cancer functions of macrophages: enhanced tumor infiltration, decreased angiogenic potential, reduced efferocytosis, and improved support of CD8+ T cell activity. Mechanistically, different AKT isoforms are involved, through both PI3K dependent and independent mechanisms. Assessment of human blood and breast tumors revealed correlations between ABCA1 in macrophages and angiogenic potential, *VEGFA*, and CD8 T cell abundance and activity, highlighting the clinical relevance of our findings. The culmination of the effects of ABCA1 on MC function were demonstrated through increased tumor growth and metastasis in mice with MC specific knockout of ABCA1. Therefore, modulating ABCA1 activity within MCs may represent a novel approach to immune therapy.

## INTRODUCTION

Immune therapies such as chimeric antigen receptor T-cell (CAR T cells) and immune checkpoint blockade (ICB) have had enormous impact on the clinical treatment of cancer, and in some cases even offering curative therapy. However, CAR T cells require a cancer specific or enriched epitope to target, restricting their broad applicability. Furthermore, ICB has limited efficacy in many solid tumors, including metastatic breast cancer. Pembrolizumab (anti programmed cell death protein 1; αPD1) in combination with nab-paclitaxel for patients with advanced triple-negative breast cancer (TNBC) positive for the PD1 ligand (PD-L1) improves pathologic complete response and event-free survival (1). However, the vast majority of patients do not respond (1–3). Although αPD1 in the neoadjuvant setting delays progression, it is not curative. Given that metastatic disease accounts for the majority of mortality associated with breast cancer, 1 in 8 women will be diagnosed with invasive breast cancer within their lifetime, and breast cancer being second leading cause of cancer-related deaths in women, it is of paramount importance to probe mechanisms that may improve the efficacy of ICB.

Cholesterol and regulators of its homeostasis have emerged as important modulators of tumor immunology (4, 5). Elevated circulating cholesterol has been associated with breast cancer recurrence (6). On the other hand, use of cholesterol lowering medication (statins) is associated with increased time to breast cancer recurrence (7–9). This association is also true for TNBC patients taking statins, demonstrating improved overall survival and breast cancer specific survival (10). However, statins have a high first-pass metabolism, allowing for tumors to escape and produce their own cholesterol (11, 12). Thus, therapeutics targeting downstream metabolism or actions of cholesterol are required to truly harness the clinical associations between cholesterol and progression.

Cholesterol has been shown to influence several different immune cell types. Since ICB already targets T cells, we have focused on myeloid immune cells, which have been strongly implicated in tumor progression, resistance to conventional therapy, and resistance to ICB (4, 13, 14). However, myeloid cells are required for antigen presentation, T cell support and effective anti-tumor responses (15). Therapeutic strategies where they are removed or inhibited have failed (16). Therefore, rather than ablating or inhibiting their activities altogether, efforts to re-educate myeloid cells towards anti-tumor phenotypes are likely to have significant clinical benefit (17–19). An oxysterol metabolite of cholesterol, 27-hydroxycholesterol, was found to work through the liver x receptors in myeloid immune cells to robustly suppress T cell proliferation and function, and stimulate the secretion of pro-tumor extracellular vesicles, ultimately promoting the growth of tumors and metastatic lesions (17, 20–23). The negative regulator of cholesterol catabolism, NR0B2, was found to suppress several aspects of the inflammasome in myeloid cells, resulting in decreased expansion of regulatory T cells (Tregs) (24, 25). A recently developed agonist of NR0B2 was demonstrated to have significant anti-tumor activity in murine models (26). Therefore, regulators of cholesterol homeostasis appear to have unique capabilities to modulate myeloid cell function, potentially ‘re-educating’ them.

The cholesterol homeostatic pathway involves several proteins, offering potentially new therapeutic targets. Here, we describe how the cholesterol efflux protein, ABCA1, alters several aspects of macrophage function. These functions have important implications within the tumor microenvironment. Correlations of ABCA1 and markers for myeloid cells and T cells in human breast tumors corroborate our preclinical findings. Collectively, our data suggest that ABCA1 represents a target to re-educate myeloid cells for the treatment of breast and other solid tumors.

## RESULTS

### The cholesterol efflux and translocation regulatory protein, ABCA1, is expressed in myeloid immune cells and is associated with increased survival

Given previous studies and clinical data supporting a role for cholesterol metabolism in influencing tumor associated myeloid cell function, we sought to identify proteins that are involved in cholesterol homeostasis that are (1) expressed in myeloid cells, and (2) associated with survival of breast cancer patients. In doing so, we identified ABCA1, a protein involved in cholesterol efflux and translocation from the inner leaflet to the outer leaflet of the plasma membrane (4, 27). Multiplex immunofluorescence (mIF) of human breast tumors found that while the majority of cells stained positive for ABCA1, CD11B+ myeloid cells displayed gradients of expression, generally higher than other cell types (**Fig. 1A**). Certain tracks of CD11B+ cells had significantly higher expression of ABCA1. This is supported by single cell RNA-seq (scRNA-seq) analysis of healthy breast tissue where macrophages had high expression (data mined from the Human Protein Atlas (28); **Fig. 1B**, corresponding cluster map in **Supplementary Fig. 1A**). scRNA-seq of human breast tumors also reveals elevated expression of ABCA1 within myeloid cells, and in particular macrophages; consistent across three different cohorts (**Fig. 1C**). Similarly, scRNA-seq of mammary tumors from genetically engineered mouse models showed higher expression of ABCA1 in monocyte/macrophage cells (**Fig. 1C**). Assessment of traditionally polarized murine bone marrow derived macrophages (BMDMs) found modest decreases in *Abca1* in M1 polarized compared to M0 or M2 (**Supplementary Fig. 1B**). Expression of *Abca1* was elevated in BMDMs compared to bone marrow derived dendritic cells (BMDCs), or CD11C+ (dendritic cell-like) or CD11B+;Ly6G+ (granulocyte) cells isolated naïve from mice. Tumor derived CD11B+ (pan-myeloid) cells had elevated *Abca1* compared to CD11B+ cells isolated from spleen or bone, or CD11C+ cells (**Supplementary Fig. 1B**). Thus, ABCA1 is expressed in MCs, including those within tumors.

**Figure 1:**
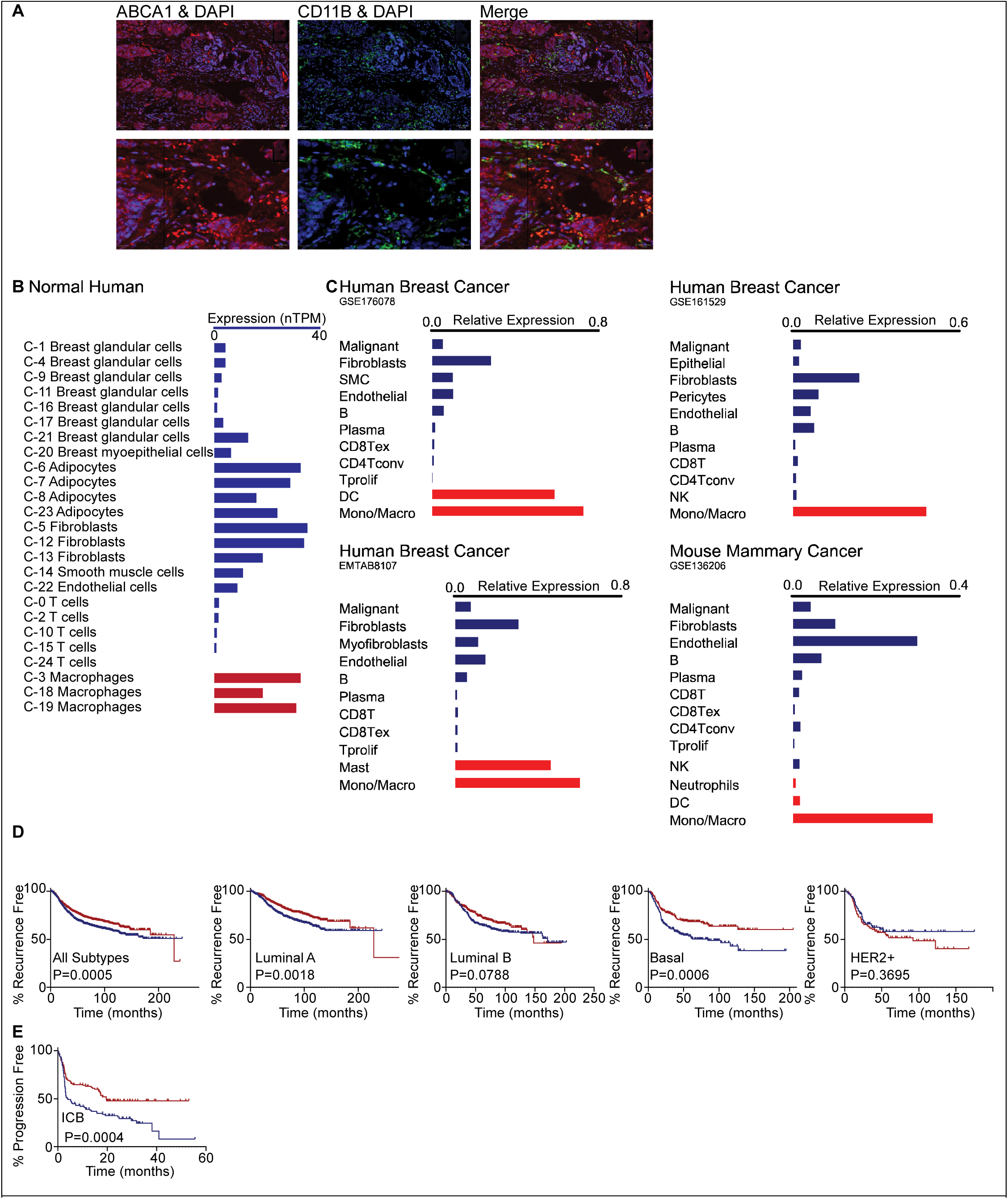
ABCA1 is expressed in breast tumor associated myeloid cells and is positively associated with increased survival. **(A)** Breast tumor sections were stained for ABCA1 (red), the pan-myeloid cell marker CD11B (green), and the nuclear stain DAPI (blue). **(B)** Assessment of ABCA1 expression by single cell RNA-seq of healthy breast tissue indicates relatively high expression in cells annotated as macrophages (data from the Human Protein Atlas; corresponding cluster map in **Supplementary Fig. 1**; proteinatlas.org (29)). Red bars are of the myeloid cell lineage. (**C**) Single cell RNA-seq of breast/mammary tumors indicates relatively high expression of ABCA1 in myeloid cells compared to other cells. Red bars are of the myeloid cell lineage. Data was obtained from the indicated databases and accessed through TISCH2 (30). GSE176078: 26 tumors, 89,471 cells. GSE161529: 52 tumors, 332,168 cells. EMTAB8107: 14 tumors, 33,043 cells. GSE136206: 27,352 cells. (D) ABCA1 expression is associated with increased survival. The Kaplan-Meier plotter was used to probe associations between ABCA1 expression and recurrence-free survival when considering all subtypes, Luminal A only, Luminal B only, Basal only, or HER2+ only (from left to right) (P values were determined using the Log-rank (Mantel-Cox) method except for Luminal B where the Gehan-Breslow-Wilcoxin test was used). (E) Expression of ABCA1 in tumors from patients treated with immune checkpoint blockers (ICB) is associated with an increased progression free survival time. Tumor types included for this analysis were bladder, esophageal adenocarcinoma, glioblastoma, hepatocellular carcinoma, head and neck squamous cell carcinoma, melanoma, nonsmall cell lung cancer and urothelial cancer. P value was determined using the Log-rank (Mantel-Cox) method. The Kaplan-Meier Plotter webtool uses aggregated data from GEO, EGA, and TCGA (31).

Importantly, elevated expression of ABCA1 in tumors is associated with increased recurrence free survival, when considering all breast cancer subtypes collectively, or when only considering Luminal A, Luminal B or Basal subtypes separately (**Fig. 1D**). Our analysis was underpowered to make firm conclusions regarding associations in HER2+ cases. Interestingly, in patients taking immune checkpoint blockade, high ABCA1 expression is also associated with prolonged progression free survival (all cancer types considered, **Fig. 1E**).

Collectively, these data suggest that ABCA1 activity plays a role in regulating myeloid cells in ways that are important for tumor progression. Therefore, we performed several experiments to assess whether ABCA1 influenced myeloid cell functions critical for shaping tumor pathophysiology, specifically (1) tumor infiltration, (2) angiogenesis, (3) phagocytosis (4) efferocytosis, and (5) support of T cell expansion/function. We focused on macrophages since they are an abundant myeloid cell type in breast tumors and metastatic lesions (15). siRNA-mediated knockdown of ABCA1 resulted in the expected decrease in ABCA1 mRNA, protein and cholesterol efflux (**Supplementary Fig. 1C-G**). While decreasing efflux, siRNA against ABCA1 also significantly increased cholesterol accumulation within the inner-leaflet of the plasma membrane (**Supplementary Fig. 1H**).

### ABCA1 increases bone marrow derived macrophage (BMDM) infiltration into tumor spheroids

In order to support immune activity, myeloid cells such as macrophages must be able to penetrate tumors. When ABCA1 was knocked down in BMDMs, they were restricted to the surface of 4T1 mammary cancer spheroids, while control siRNA treated BMDMs were able to infiltrate (**Fig. 2A**). This was confirmed using a second siRNA targeting ABCA1 and in a separate experiment where BMDM infiltration was also assessed by flow cytometry of dissociated spheroids (**Fig. 2B**). BMDMs treated with PSC833, a small molecule reported to inhibit ABCA1 efflux (32), also demonstrated reduced infiltration (**Fig. 2C**). Conversely, ABCA1 overexpression increased the capacity to infiltrate 4T1 tumor spheroids (**Fig. 2D**). Similar results were obtained when assessing infiltration into E0771 mammary cancer spheroids (**Fig. 2E-H**). Therefore, data generated using multiple approaches to alter ABCA1 and two different spheroid types strongly indicates that ABCA1 activity results in increased ability of macrophage entry into the tumor microenvironment.

**Figure 2:**
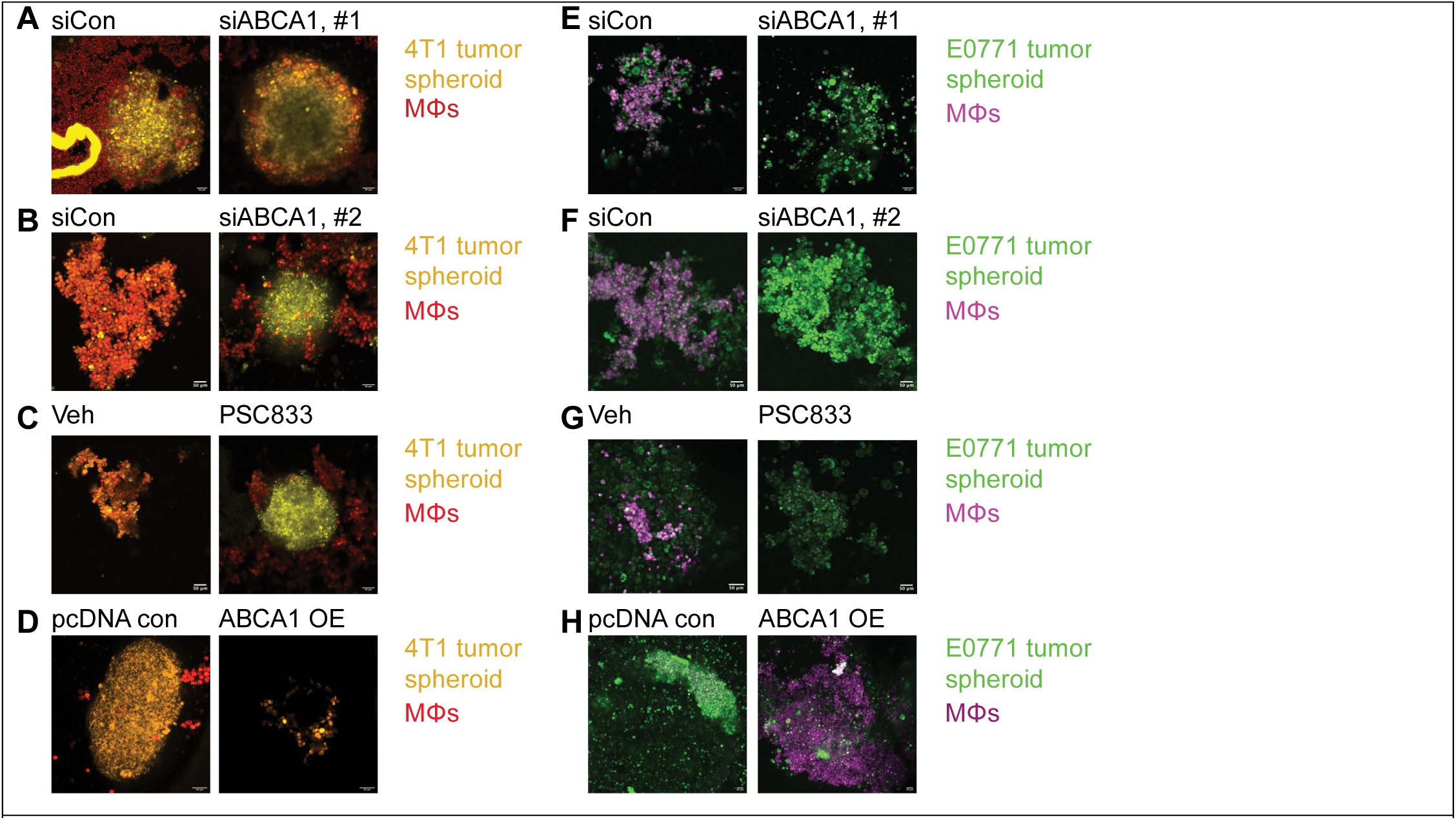
ABCA1 activity in macrophages increases their capacity to infiltrate mammary cancer spheroids. **(A)** BMDMs (MΦs) were transfected with control or siRNA against ABCA1 for 48h prior to being stained with Cell Trace Red and overlayed onto 4T1 tumor spheroids previously stained with HCS Nuclear Mask and Cell Mask Orange, in a ultra-low attachment dish. Confocal microscopy was performed on a Ziess LSM 900 using airyscan. Z stack images were obtained and analyzed using Image J. Maximum intensity z projections were created for representative confocal images. **(B)** A second siRNA targeting a different region of *Abca1* in MΦs also resulted in decreased infiltration of 4T1 tumor spheroids. **(C)** A small molecule inhibitor of ABCA1, PSC833, decreased the ability of MΦs to infiltrate 4T1 tumor spheroids. BMDMs were treated with 1µM PSC833 for 24h. BMDMs were washed and stained prior to overlay onto 4T1 spheroids. **(D)** Overexpression of ABCA1 in MΦs increased their capacity to infiltrate 4T1 tumor spheroids compared to empty vector (pcDNA) control transfected MΦs. **(E-H)** Manipulating ABCA1 had similar effects on the ability of MΦs to infiltrate E0771 tumor spheroids.

### ABCA1 in BMDMs reduces angiogenesis (endothelial cell tube formation)

Promotion of angiogenesis is a key attribute of tumor associated macrophages and is generally seen as pro-tumor. BMDMs were cocultured above HUVEC cells (human umbilical vein endothelial cells) growing in a basement membrane, separated by a 0.4µm porous membrane. siRNA mediated knockout of ABCA1 in BMDMs resulted in significantly increased tube formation (**Fig. 3A**). Pretreatment with the ABCA1-efflux inhibitor PSC833 similarly enhanced tube formation although more subtly than knockdown (**Supplementary Fig. 3**). On the other hand, overexpression of ABCA1 in BMDMs led to near-complete inhibition of tube formation (**Fig. 3B**). Thus, ABCA1 within macrophages significantly impairs their ability to support angiogenesis.

**Figure 3:**
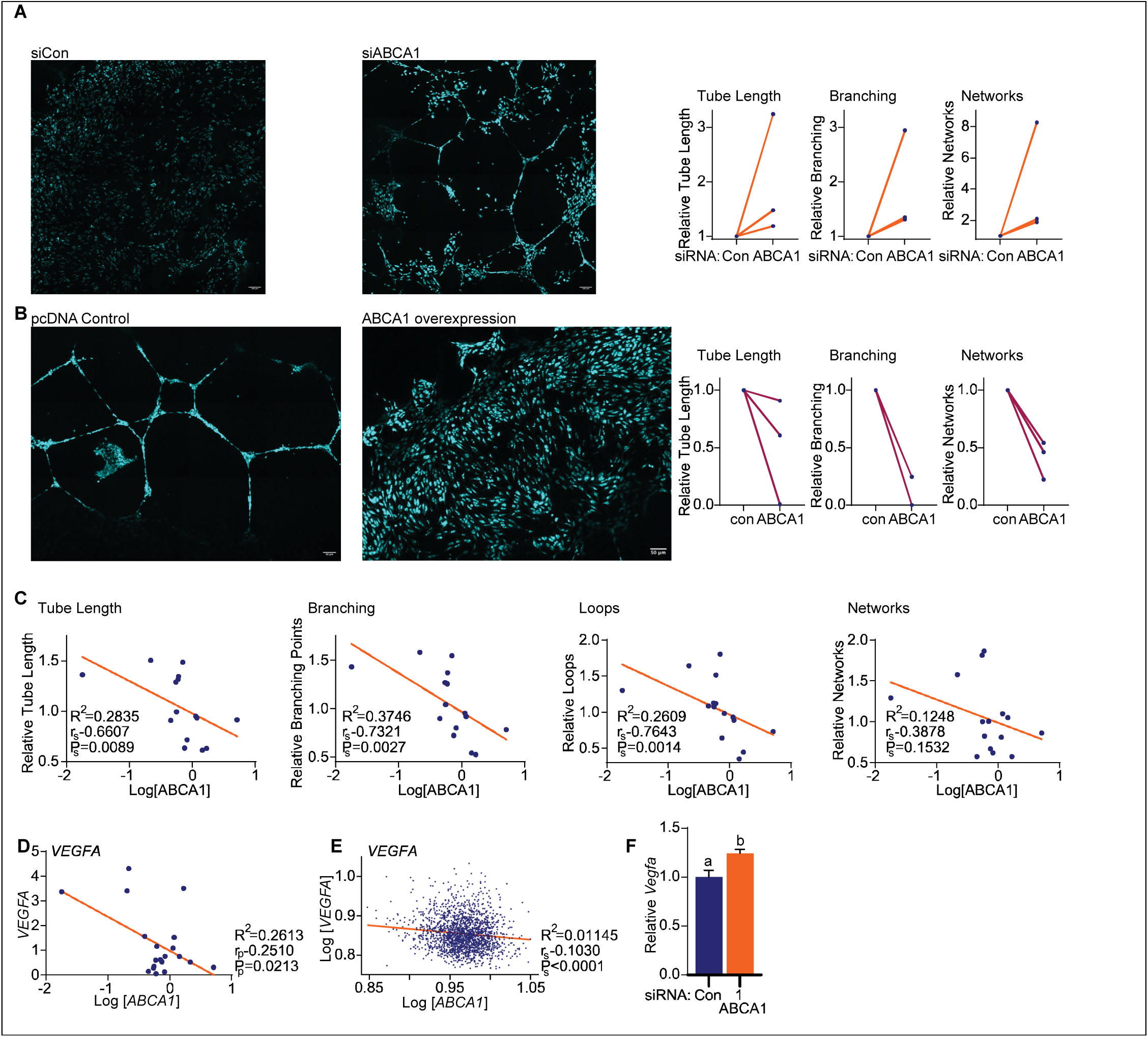
ABCA1 activity in macrophages decreases their ability to promote neo-angiogenesis. **(A)** siRNA against ABCA1 in MΦs results in increased tube formation by HUVEC cells. MΦs were transfected with control or siRNA against ABCA1 for 48h before being placed on top of HUVEC cells embedded in Geltrex. Macrophages and HUVECs were separated by a 0.4µm Boyden Chamber only allowing for exchange of soluble factors. 24h later, HUVECs were stained with Calcein AM to visualize Live Cells. Confocal microscopy was performed using Ziess LSM900, while performing tile scans and z stacks. Tube formation metrics were quantified by Wimasis. Representative images to the left of quantified data for 3 independent experiments normalized to control for batch effects. Total tube length, branching and networks were quantified **(B)** ABCA1 overexpression in MΦs results in decreased tube formation. Same experimental setup as (A), but MΦs were transfected with either control (pcDNA) or an ABCA1 overexpression plasmid. **(C)** Human PBMCs were sourced from the whole blood of females with no known cancer, and differentiated into macrophages. Media from 48h of conditioning was added to pre-seeded HUVECS and confocal microscopy was performed as described above. Linear regression was performed to evaluate potential correlations between *ABCA1* expression and different aspects of tube formation (length, branching, loops and networks). Spearman’s rank correlation (r_s_) was determined and corresponding P (P_s_) value reported. **(D)** Human PBMCs were probed for the expression of *VEGFA*, and linear regression performed comparing *ABCA1* to *VEGFA*. The Pearson correlation coefficient (r_p_) and corresponding P (P_p_) value reported. **(E)** Linear regression comparing expression of *ABCA1* to *VEGFA* in human breast tumors. Data obtained from the TCGA (Breast Invasive Carcinoma). Spearman’s rank correlation (r_s_) was determined and corresponding P (P_s_) value reported. **(F)** *Vegfa* mRNA expression is increased in murine MΦs transfected with siRNA against ABCA1. Different letters denote statistical significance (P<0.05).

To explore this in humans, peripheral blood mononuclear cells (PBMCs) were isolated from female volunteers with no known cancer, and evaluated for *ABCA1* expression and their capacity to stimulate tube formation. We observed an inverse correlation between PBMC-derived macrophage *ABCA1* expression and their ability to stimulate tube formation, as measured by tube length, branching and loops (**Fig. 3C**). Since our system prevented direct interaction between macrophages/PBMCs and endothelial cells, it can be concluded that a soluble factor was mediating the effects on tube formation. Tube formation and angiogenesis are modulated by several factors, a predominant one being VEGFA. Interestingly, *ABCA1* mRNA expression in a different cohort of human PBMCs was also inversely correlated with *VEGFA* (Spearman r of -0.251, P=0.0213; **Fig. 3D**). Analysis of breast tumor samples also revealed a negative correlation between *ABCA1* and *VEGFA* (TCGA analysis, Spearman r of -0.1030, P<0.0001; **Fig. 3E**). Furthermore, murine BMDMs transfected with siRNA against ABCA1 had slightly increased *Vegfa* (**Fig. 3F**; data confirmed by RNAseq, **Fig. 7**). Therefore, some as yet unknown factor downstream of ABCA1 activity influences neo-angiogenesis, potentially through a decrease in VEGFA.

### ABCA1 decreases efferocytosis but not phagocytosis in BMDMs

It has been reported that macrophages can directly kill cancer cells (33, 34). Related and perhaps more well appreciated is their ability to phagocytose cancer cells or their debris, resulting in presentation of neoantigen and subsequent activation of T cells. Macrophages also engulf particles, especially apoptotic bodies, through a specialized type of phagocytosis, termed efferocytosis. Efferocytosis does not culminate in antigen presentation and instead evokes a highly immune suppressive macrophage phenotype (35, 36). Indeed, macrophage cell therapy has faced a significant hurdle in that macrophages preferentially efferocytose cancer cells, and significant efforts are being made to inhibit efferocytosis therapeutically (37, 38).

siRNA mediated knockdown of ABCA1, treatment with PSC833 or overexpression had no measurable effects on phagocytosis of inactivated, E. coli conjugated beads (**Fig. 4A-C**). However, the relative efferocytosis of apoptotic neutrophils by macrophages was enhanced by two different siRNAs against ABCA1 (**Fig. 4D-E**). Likewise, the ABCA1 inhibitor PSC833 also increased efferocytosis (**Fig. 4F**). Conversely, ABCA1 overexpression reduced efferocytosis (**Fig. 4G**). The conventional approach to assessing efferocytosis utilizes apoptotic neutrophils as “bait”. More relevant to cancer biology, we also found that siRNA knockdown of ABCA1 increased efferocytosis of apoptotic E0771 cancer cells, as did PSC833 (**Fig. 4H-I**). Overexpression of ABCA1 significantly decreased the efferocytosis of cancer cells (**Fig. 4J**). Therefore, ABCA1 does not influence total phagocytic activity but does decrease efferocytosis, so there is likely a net increase in immunogenic phagocytosis (non-efferocytotic phagocytosis).

**Figure 4:**
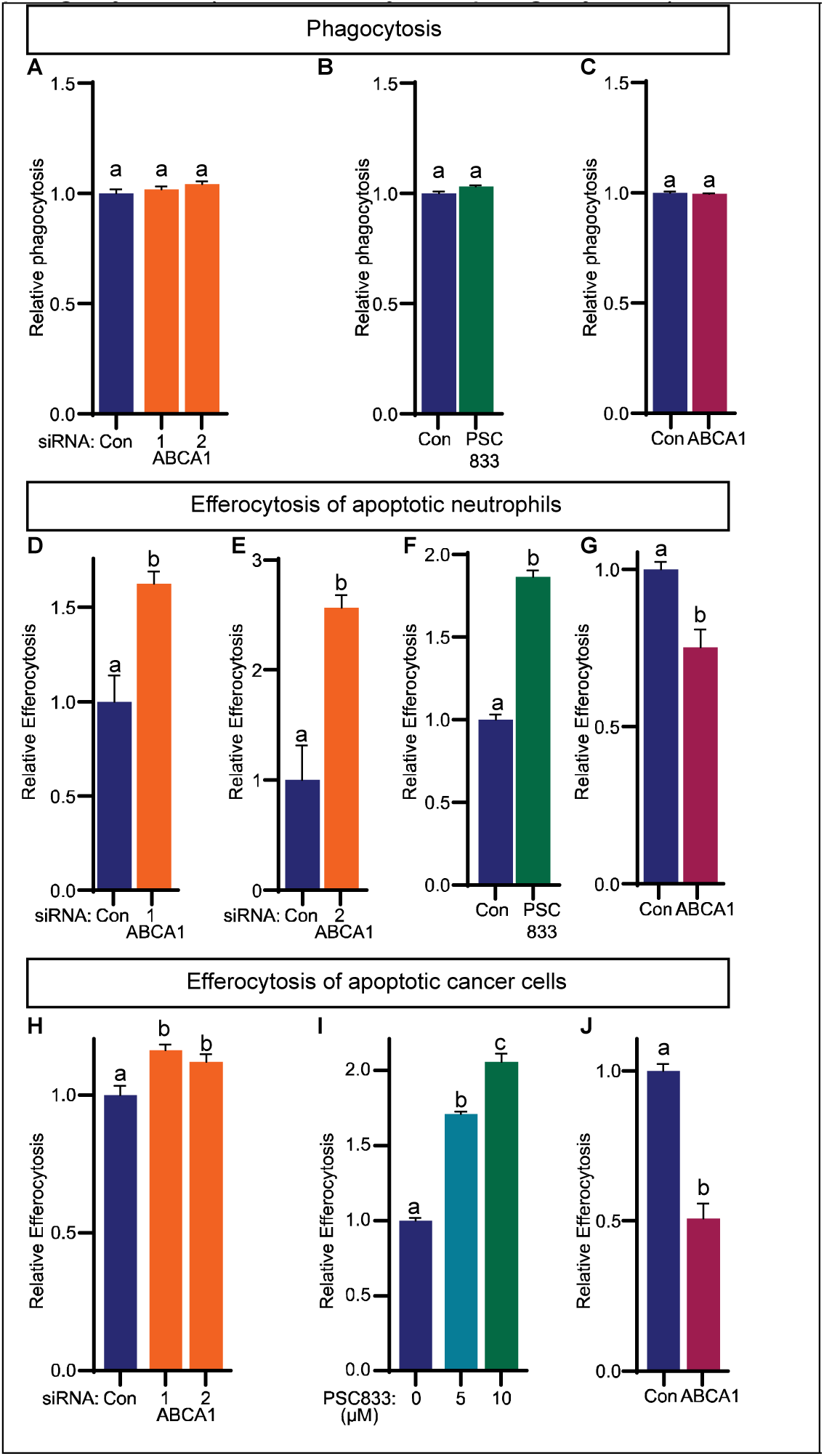
ABCA1 overexpression in BMDMs results in decreased efferocytosis, while loss of ABCA1 increases efferocytosis. **(A-C)** No difference in phagocytosis was observed between control BMDMs or those transfected with two different siRNAs against ABCA1, treated with the ABCA1 inhibitor PSC833 nor ABCA1 overexpression. **(D-E**) two independent siRNAs against ABCA1 resulted in increased efferocytosis of apoptotic neutrophils. **(F)** Likewise, treatment of BMDMs with PSC833 also increased efferocytosis. **(G)** Overexpression of ABCA1 decreased efferocytosis. **(H-J)** Same as for (D-G) but the efferocytosis ‘bait’ was apoptotic E0771 cells instead of neutrophils.

### ABCA1 in BMDMs supports T cell migration, expansion and anti-cancer function

In order to mount a successful anti-tumor immune response, myeloid cells must effectively support T cells. The first step is T cell recruitment to tumors; T cell exclusion from tumors being considered a major barrier to ICB. BMDMs were transfected and cultured in the presence of a peptide fragment of ovalbumin (OVA) and activated with LPS. T cells from OTI mice have been engineered to express a transgenic T-cell receptor designed to recognize immunogenic residues within OVA in the context of major histocompatibility class I molecules (H2-Kb). Pan-OTI T cells were then co-cultured with the BMDMs for 72hrs. The T cells were then labeled with a stain (carboxyfluorescein succinimidyl ester; CFSE) and placed in the top well of a Boyden chamber with OVA expressing E0771 mammary cancer cells (E0771-OVA) in the lower well (experimental setup depicted in **Figure 5A**, left panel). Migration through the transmembrane towards the E0771-OVA cells was assessed 10hrs later.

**Figure 5:**
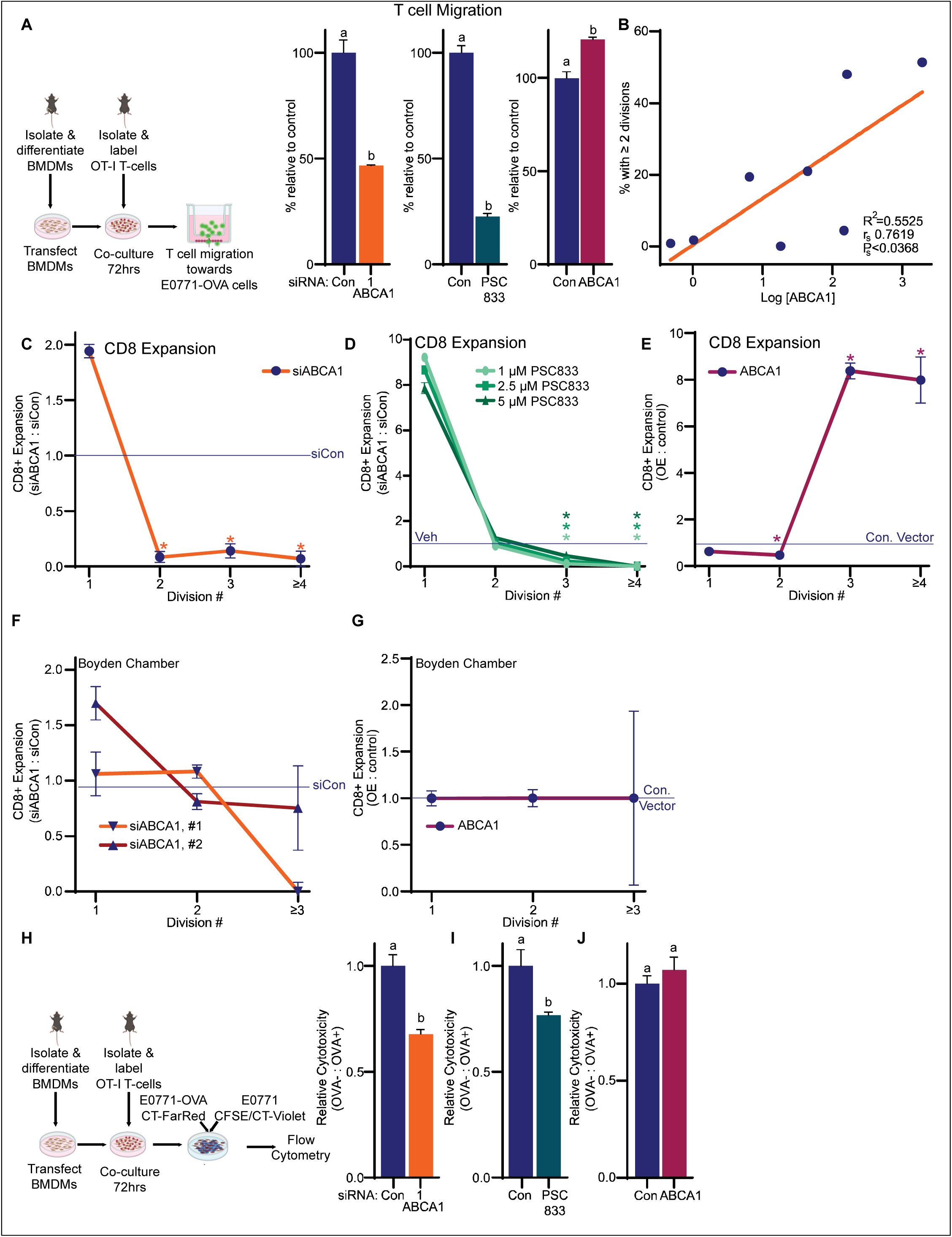
ABCA1 activity results in BMDMs that support T cell migration, expansion and anti-cancer cytotoxic function. **(A)** BMDMs were transfected and cultured in the presence of OVA and LPS. Pan-OTI T cells were co-cultured with the BMDMs for 72hrs. The T cells were then labeled with CFSE and placed in the top well of a Boyden chamber and allowed to migrate towards E0771-OVA mammary cancer cells. Cartoon of experimental approach to the left of quantified data. **(B)** Human, female PBMCs were isolated from whole blood and bead-sorted for monocytes and T cells. Monocytes were differentiated into macrophages prior to assessment of *ABCA1* expression by qPCR and culture with activated T cells from the same donor. T cell proliferation was determined by dilution of CFSE stain, by flow cytometry. Linear regression was performed comparing *ABCA1* expression to the percentage of T cells that had undergone more than 2 divisions. Spearman’s rank correlation (r_s_) was determined and corresponding P (P_s_) value reported. **(C)** CD8+ T cell expansion is decreased when activated T cells are co-cultured with BMDMs transfected with siRNA against ABCA1. A ratio between the proportion of T cells expanded under siABCA1 conditions to siControl conditions is shown, with the blue line being the siControl conditions. The percentage in each clonal generation shown in **Supplementary Fig. 4A**. A second siRNA against ABCA1 yielded similar results (**Supplementary Fig. 4B**) **(D)** CD8+ T cell expansion is decreased when activated T cells are co-cultured with BMDMs pretreated with the ABCA1 inhibitor, PSC833. BMDMs were pre-treated with indicated doses of PSC833 for 24hr prior to co-culture with activated T cells. A ratio between the proportion of T cells expanded under treatment conditions to vehicle control conditions is shown, with the blue line being the vehicle conditions. The percentage in each clonal generation shown in **Supplementary Fig. 4D**. **(E)** CD8+ T cell expansion is increased when activated T cells are co-cultured with BMDMs transfected with an expression plasmid for ABCA1. A ratio between the proportion of T cells expanded under ABCA1 overexpression conditions to control vector conditions is shown, with the blue line being the control conditions. The percentage in each clonal generation shown in **Supplementary Fig. 4E**. **(F)** The robust effects of siABCA1 in BMDMs on CD8+ T cell expansion are not as apparent when the BMDMs are separated from T cells in a Boyden chamber. The percentage in each clonal generation shown in **Supplementary Fig. 4I. (G)** The robust effects of overexpressing ABCA1 in BMDMs on CD8 T cell expansion are lost when the BMDMs are separated from T cells in a Boyden chamber. The percentage in each clonal generation shown in **Supplementary Fig. 4J**). **(H)** Anti-cancer cytotoxicity of T cells is influenced after coculture with BMDMs with altered ABCA1. BMDMs were transfected with siControl or siRNA against ABCA1 for 48h. They were then treated with OVA prior to co-culture with OT-I T cells for 72h. T cells were counted and an equal number were co-cultured with E0771 and E0771-OVA cells at a starting ratio of 1:1, each stained with a different color. After 24hrs, remaining E0771 and E0771-OVA cells were quantified by flow cytometry. The experimental setup is illustrated to the left. Data is displayed as a ratio of OVA-to OVA+ E0771 cells, so decreases are a reflection of antigen-specific, T cell-mediated cytotoxicity. **(I)** Treatment of BMDMs with the ABCA1 inhibitor PSC833 results in T cells with decreased cytotoxic activity. Experimental setup and details as in H. **(J)** Overexpression of ABCA1 in BMDMs did not robustly influence T cell cytotoxic capacity against E0771-OVA cells. Experimental setup and details as in H. Asterisks (*) or different letters denote statistical significances (P<0.05). A, H, I & J: Student’s T Test. C-G: Student’s T Test for each division, comparing treated to control condition. Cartoons generated and adapted from BioRender.

siRNA knockdown of ABCA1 in BMDMs resulted in decreased T cell migration towards cancer cells, as did treatment of BMDMs with PSC833 (**Fig. 5A**, right panel). Overexpression of ABCA1 resulted in increased T cell migration towards E0771-OVA cells (**Fig. 5A**). Therefore, ABCA1 within macrophages results in increased T cell migration, an important finding given that T cell restriction from tumors is considered a major obstacle to immune therapies.

When T cells are activated, they undergo a rapid clonal expansion, an essential component of the acquired immune response. To explore the potential relationship between macrophage-ABCA1 and their ability to support T cell expansion, we acquired human PBMCs from females with no known cancer, isolated monocytes and cultured them into macrophages. T cells were isolated from the same donors and cryo-preserved prior to re-culture, activation and subsequent co-culture with the differentiated macrophages. Making use of the natural heterogeneity of *ABCA1* expression in PBMC-derived macrophages from different subjects, we observed a robust correlation between macrophage *ABCA1* gene expression and their ability to support T cell expansion (Spearman rank coefficient r_s_ of 0.7619, P=0.0368; **Fig. 5B**). This suggests that ABCA1 within macrophages shifts them into a state whereby they facilitate T cell expansion.

To test this hypothesis directly, we made use of murine BMDMs and T cells from syngeneic mice. When T cells are activated by anti-CD3/28 and then co-cultured with BMDMs pretreated with siRNA against ABCA1, there is a significant reduction in the expansion of CD8+ (cytotoxic) T cells (ratio of two conditions shown in **Fig. 5C**; percentage in each clonal generation shown in **Supplementary Fig. 4A**). A second siRNA targeting a different region of *Abca1* yielded similar results (**supplementary Fig. 4B**). Interestingly, the effects on T cell expansion were restricted to CD8+ T cells as CD4+ T cell expansion was not significantly affected (**Supplementary Fig. 4C**). Similar to knockdown, pretreatment of BMDMs with the ABCA1 inhibitor PSC833 resulted in decreased CD8+ T cell expansion, even at the lowest dose tested (1µM, **Fig. 5D** and **Supplementary Fig. 4D**). Conversely, overexpression of ABCA1 enhanced CD8+ T cell expansion (**Fig. 5E** and **Supplementary Fig. 4E**). Of note, the expansion differences when ABCA1 was knocked out, inhibited or overexpressed were most apparent in later generations (**Fig. 5** and **Supplementary Fig. 4**). This is an important finding as T cell exhaustion is considered one mechanism of resistance to immune therapies.

Treatment of BMDMs with cholesterol is known to impair their support of T cell expansion, in part through conversion to the immune-suppressive 27-hydroxycholesterol and activation of the LXRs (22, 23). As expected, supplementation of BMDMs to exogenous cholesterol resulted in decreased CD8+ T cell expansion. However, ABCA1 overexpression was able to attenuate this (**Supplementary Fig. 4F**). This was evident whether the cholesterol was delivered dissolved in DMSO or carried with cyclodextrin (**Supplementary Fig. 4F).**

The strong effects on CD8+ T cell expansion with no significant effects on CD4+ cells was of interest. In order to assess whether the pro-expansion effects of macrophage-ABCA1 on CD8+ T cells required input from helper T cells, we repeated our experiments but using bead-sorted CD8+ T cells only (rather than pan-T cells). Intriguingly, in the absence of CD4+ cells, the effects of siRNA knockdown of ABCA1 in BMDMs on CD8+ T cell expansion were muted (**Supplementary Fig. 4G**). However, ABCA1 overexpression still had some capacity to increase CD8+ T cell expansion, albeit somewhat attenuated compared to when pan-T cells were cocultured (**Supplementary Fig. 4H**). These data suggest that helper CD4+ cells are required for the full effects of BMDM-ABCA1 on CD8+ T cell expansion, even when their own expansion is not influenced.

To evaluate whether the effects of macrophage-ABCA1 on T cell expansion required direct contact, we made use of a Boyden chamber trans-well system and BMDMs from mice lacking ABCA1 in the myeloid cell lineage. Although some inhibition remained, the robust decreases in expansion observed after siRNA against ABCA1, were lost when contact between BMDMs and T cells was prohibited (**Fig. 5F, Supplementary Fig. 4I).** More strikingly, the effects of ABCA1 overexpression were lost in this Boyden chamber setup (**Fig. 5G, Supplementary Fig. 4J**). Thus, although there may be support from soluble factors, direct contact between BMDMs and T cells is required for the full effects of ABCA1.

Irrespective of expansion, the cytotoxic activity of CD8+ T cells is what governs their ultimate anti-cancer properties. To test this, T cells from OTI mice were co-cultured with BMDMs previously incubated with OVA and LPS. The T cells were then co-cultured with E0771 and E0771-OVA cancer cells for 16-24 hours. E0771 cells were labeled with CFSE or Cell Trace Violet, and E0771-OVA cells with Cell Trace Red, allowing for antigen-specific cytotoxicity to be measured (ratio of E0771-OVA : E0771) (22, 25, 26) (experimental setup outlined in **Fig. 5H**, left panel). siRNA against ABCA1 or treatment of BMDMs with PSC833 resulted in significantly reduced killing of E0771-OVA cells (**Fig. 5H-I**). Overexpression of ABCA1 in MCs did not significantly alter cytotoxicity of expanded T cells (**Fig. 5J**), which may indicate that they are already maximally functional, or potentially that overexpression in macrophages did not sufficiently increase cholesterol efflux as there would be minimal acceptor proteins in conventional culture media. Regardless, when both expansion and activity are considered, ABCA1 activity in MCs results in better support of CD8+ T cell function.

### ABCA1 expression is correlated with tumor infiltrating CD8+ T cells and markers of T cell function

Using transcriptome data from the METABRIC dataset accessed through cBioPortal, we found robust correlations between *ABCA1*, and *CD8A* or *CD8B* expression in human breast tumors (**Fig. 6A-B**), supporting our findings that ABCA1 resulted in increased CD8+ T cell migration and expansion. siRNA knockdown of ABCA1 in BMDMs followed by co-culture with activated T cells resulted in decreased expression genes associated with cytotoxic T cell activation or function: *Ifnγ*, G*zmb* and *Prf1* (**Fig. 6C-E**). Supporting these preclinical results are correlations observed in human breast tumors between *ABCA1* with *IFNγ*, *GZMB* and *PRF1* (**Fig. 6F-H**). These trends are likely driven by ABCA1 expression extrinsic to T cells (ie: in MCs), as cholesterol efflux has been shown to impair T cell expansion and activity (39, 40).

**Figure 6:**
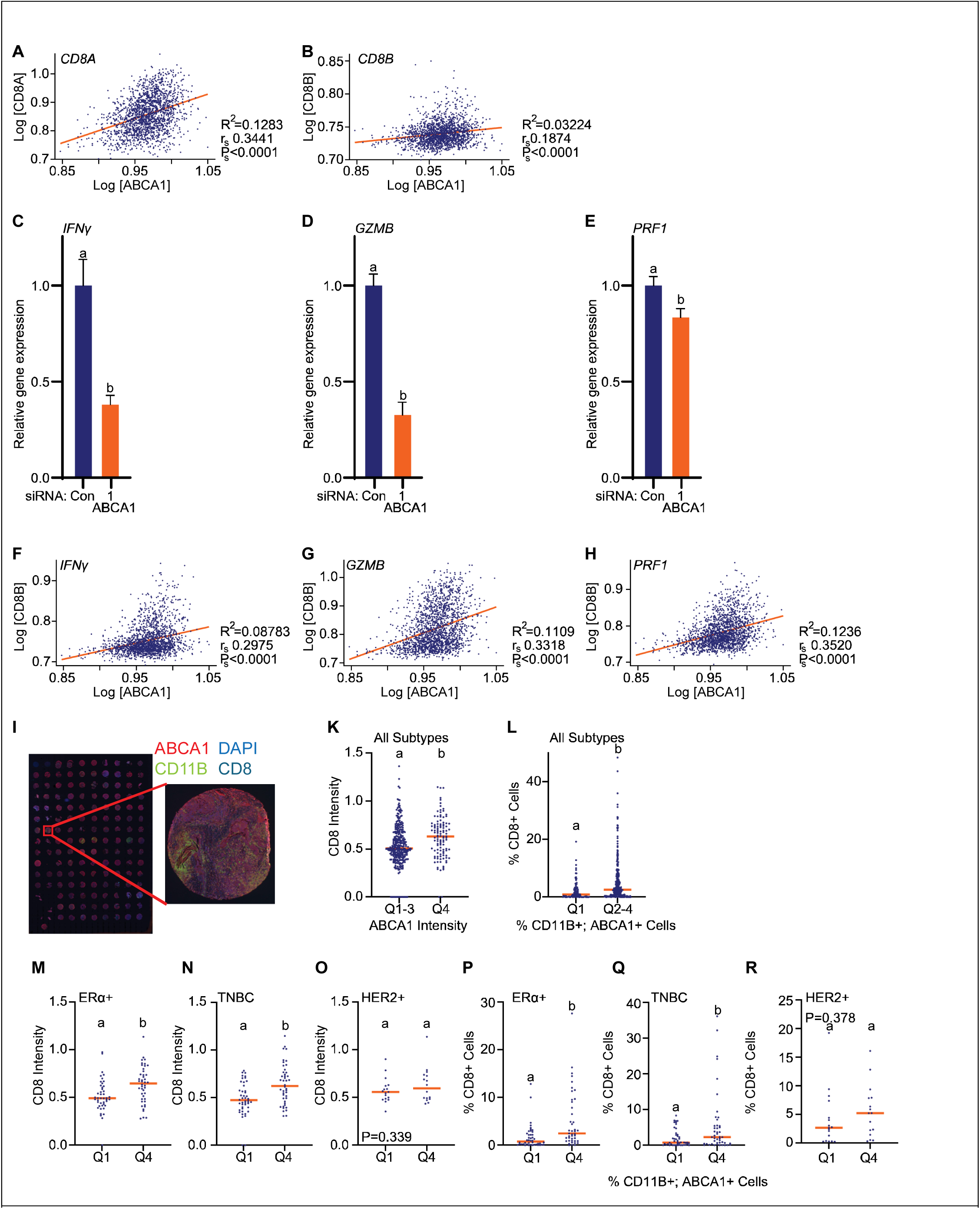
ABCA1 is correlated with markers of CD8+ T cell number and function in human breast tumors. **(A-B)** mRNA expressions for the markers for cytotoxic T cells, *CD8A* and *CD8B,* are positively correlated to *ABCA1* expression in human breast tumors. Linear regression was used to fit the line. Data obtained from the METABRIC dataset accessed through cBioPortal. Spearman’s rank correlation (r_p_) was determined and corresponding P (P_s_) value reported. **(C-E)** siRNA knockdown of ABCA1 in murine BMDMs followed by co-culture with activated T cells resulted in decreased expression genes associated with cytotoxic T cell activation or function: *Ifnγ*, G*zmb* and *Prf1*. Different letters denote statistical significance (P<0.05, Student’s T Test). **(F-H)** mRNA expression of *IFNγ, GZMB* and *PRF1*, markers of cytotoxic T cell activation/activity, were positively correlated with *ABCA1* expression. Data obtained from the METABRIC dataset accessed through cBioPortal. r_p_ and P_s_ reported. **(I-R**) Human breast tumor tissue microarrays (TMAs) were co-stained for ABCA1, the pan-myeloid cell marker CD11B, the cytotoxic T cell marker CD8, and the nuclear stain DAPI, in a mIF assay. Staining intensity as well as % cells positive for each stain was quantified. **(I)** Representative TMA image shown. **(K)** There is a positive correlation between ABCA1 and CD8 staining intensity (r_s_ =0.1853, P_s_<0.0001, data shown in **Supplementary Fig. 5B&D**). Here, data is parsed based on ABCA1 expression into the lower three quartiles and higher quartile, indicating that the mean CD8 intensity was higher in the highest ABCA1 quartile (different letters indicate statistical significance; P<0.05, Student’s T Test). **(L)** There is a positive correlation between the percentage of CD11B+ cells also expressing ABCA1, and the percentage of CD8 positive cells (r_s_ =0.2183, P_s_<0.0001, data shown in **Supplementary Fig. 5C&E**). Here, data is parsed based on ABCA1 expression into the lowest quartile and highest three quartiles, indicating that the percentage of CD8 positive cells was higher in the highest ABCA1 quartiles (different letters indicate statistical significance; P<0.05, Mann Whitney test). **(M-O)** ABCA1 staining intensities were parsed into the lowest and highest quartile and compared to CD8 staining intensity, where different breast cancer subtypes were assessed independently: ERα+, TNBC or HER2+. Significant differences were found for ERα+ (r_2_=0.1843 P_s_=0.0144) and TNBC cases (r_s_=0.2186, P_s_=0.0034), while this analysis was underpowered for HER2+ cases (r_p_=0.1400, P_p_=0.2586). different letters denote statistical significance when comparing these parsed groups (P<0.05, Student’s T Test). Corresponding correlational data can be found in **Supplementary Fig. 5F-K**. **(P-R)** The percentage of cells co-staining for CD11B and ABCA1 were parsed into the lowest and highest quartile and compared to the percentage of CD8+ cells, where different breast cancer subtypes were assessed independently: ERα+, TNBC or HER2+. Significant differences were found for ERα+ (r_2_=0.3187, P_s_<0.0001) and TNBC cases (r_s_=0.2628, P_s_=0.0003), while this analysis was underpowered for HER2+ cases (r_s_=0.1351, P_s_=0.2831). Different letters denote statistical significance when comparing these parsed groups (P<0.05, Mann Whitney test except for R where a Students T test was used). Corresponding correlational data can be found in **Supplementary Fig. 5L-Q.**

These data based on mRNA expression highlight the relevance of our findings to human disease. To more definitively ascribe these correlations to myeloid cell expression of ABCA1, and assess these correlations at the protein level, we developed a mIF assay. Our mIF assay was optimized to simultaneously probe tumor microarrays (TMAs) for (i) CD11B, a pan-MC marker, (ii) CD8, a marker of cytotoxic T cells, and (iii) ABCA1. Five unique breast cancer TMAs, procured from different sources, were stained and assessed (representative image in **Fig. 6I**). Three TMAs included normal or adjacent normal tissue, however, we were insufficiently powered to directly compare ABCA1 staining intensity of these cases to malignant (**Supplementary Fig. 5A**). Importantly, we found that ABCA1 staining intensity was positively correlated with CD8 staining intensity, when considering malignant samples only. Although a correlation was present when ABCA1 and CD8 were considered as continuous variables (**Supplementary Fig. 5B & D**), a non-linear relationship was observed, so we parsed intensity by quartile (**Fig. 6K**). Likewise, a correlation was found between the percentage of CD11B+ cells also expressing ABCA1 and the percentage of CD8+ cells (**Supplementary Fig. 5C & E**); tumors with the highest three quartiles of CD11B+, ABCA1+ double positive cells were associated with the highest percentage of CD8+ cells (**Fig. 6L**). This suggests that there may be a threshold effect for ABCA1, where a minimum amount is most associated with cytotoxic T cell abundance. ABCA1 staining intensity was also positively associated with CD8 when considering only ERα+ or TNBC tumors (**Fig. 6M-R**, **Supplementary Fig 5 F-Q**). We did not have sufficient power to make conclusions regarding HER2+ tumors.

Therefore, our findings in murine models that ABCA1 within myeloid cells alter CD8 T cell expansion and activity are supported by data from human tumors, suggesting conserved biology between the two species and that this is a relevant axis in humans.

### Loss of ABCA1 in macrophages is associated with altered phosphoproteome and transcriptome enriched for growth factor, membrane initiated signaling and AKT signaling

In order to understand the mechanism by which ABCA1 alters macrophage function, we performed RNA-seq, ATAC-seq and phosphoproteomics. RNA-seq analysis revealed 4901 differentially expressed genes (DEGs) between control and siRNA ABCA1 knockdown macrophages (FDR of <0.05, **Fig. 7A**). Enrichment of genes was observed in gene ontology terms associated with functional processes that we had previously identified (motility & taxis, angiogenesis, and cell to cell communication; **Supplementary Fig. 6**): (1) differentiation; as in, differentiation into different functional state or polarity, (2) motility and taxis, a likely reflection of altered ability to infiltrate tumor spheroids. (3) angiogenesis and tube formation, and (4) cell to cell signaling, a likely reflection of altered T cell expansion and function. Interestingly, several GO terms involving response to growth factors and receptor mediated signaling were also enriched for (**Supplementary Fig. 6**). Gene ontology analysis revealed enrichment of up-regulated genes in pathways such as cancer and cancer-metabolism (**Fig. 7B**). Classic macrophage polarization states of M1 or M2 were not significantly enriched for, suggesting that loss of ABCA1 re-educated macrophages into unique functional states.

**Figure 7:**
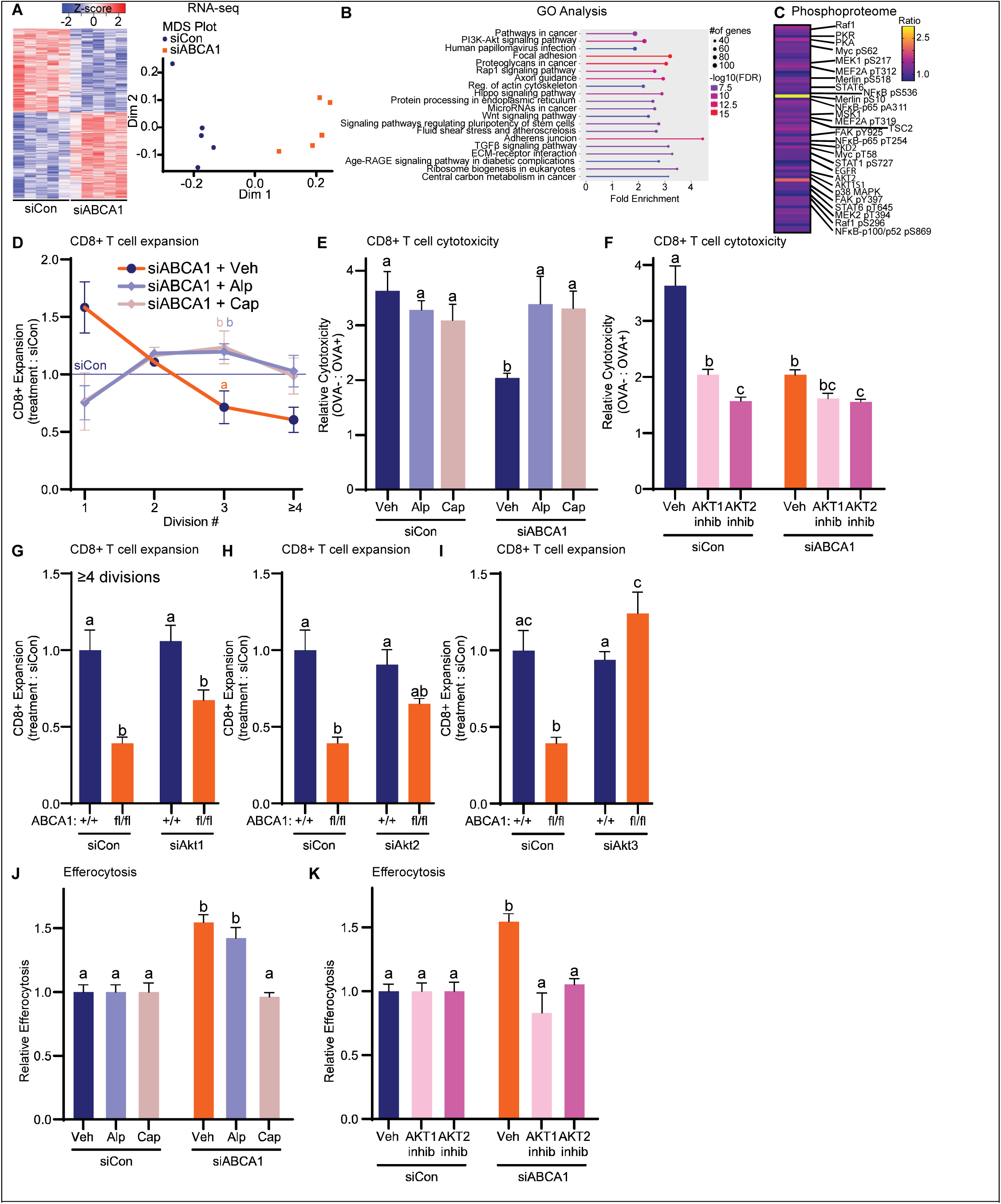
Loss of ABCA1 in macrophages associated changes in gene expression and chromatin accessibility, and an altered phosphoproteome, implicating PI3K and AKT signaling. **(A)** RNA-seq and ATAC-seq was performed on BMDMs that had been transfected with control or siRNA against ABCA1. A heatmap of differentially expressed genes (DEGs) is displayed to the left of an MDS plot indicating separation between the groups in two dimensions of transcriptional space. **(B)** Gene ontology (GO) analysis of pathways enriched after knockdown of ABCA1, indicating increased enrichment for genes associated with membrane initiated signaling such as the PI3K-AKT pathway. Further GO and KEGG pathway analyses are shown in **Supplementary Figs. 6 & 7**. ATAC-seq analyses are shown in **Supplementary Figs. 8-10**. **(C)** A phospho-protein array compared control versus siABCA1. Significantly different ratios between the phospho-protein and parental protein shown as a heatmap. **(D)** Treatment of BMDMs with alpelisib (PI3K inhibitor) or capivasertib (pan-AKT inhibitor) attenuated the subsequent inhibition of CD8+ T cell expansion when ABCA1 was knocked out. Different letters denote statistically significant differences for division 3. **(E)** Treatment of BMDMs with alpelisib or capivasertib attenuated the subsequent inhibition of anti-cancer cytotoxic T cell activity when ABCA1 was knocked out. Experimental setup outlined in Fig. 5H. Different letters denote statistical significance (P<0.05, 1-Way ANOVA followed by Šidák’s posthoc). **(F)** Neither treatment of BMDMs with an AKT1-selective inhibitor (A-674563) or and AKT2-selective inhibitor (CCT128930) were able to reverse the effects of siRNA-mediated loss of ABCA1 on anticancer cytotoxic T cell activity. Experimental setup outlined in Fig. 5H. Different letters denote statistical significance (P<0.05, 1-Way ANOVA followed by Šidák’s posthoc). **(G-I)** Anti-cancer cytotoxic T cell activity after co-culture with BMDMs from control ABCA1+/+;LysMCre+ mice or BMDMs lacking ABCA1 (from ABCA1fl/fl;LysMCre+ mice), transfected with control or siRNA against AKT1, AKT2 or AKT3. Experimental setup outlined in Fig. 5H. Different letters denote statistical significance (P<0.05, 1-Way ANOVA followed by Šidák’s posthoc). **(J)** Efferocytosis by BMDMs treated with alpelisib or capivasertib. Different letters denote statistical significance (P<0.05, 1-Way ANOVA followed by Šidák’s posthoc). **(K)** Efferocytosis by BMDMs treated with A-674563 or CCT128930 (AKT1 or 2 inhibitors respectively). Different letters denote statistical significance (P<0.05, 1-Way ANOVA followed by Šidák’s posthoc).

We observed enrichment in the PI3K-Akt signaling pathway as well as the Rap1 signaling pathway which has crosstalk with the PI3K-Akt pathway (**Fig. 7B**) (41). Closer investigation revealed the expression of several genes within the PI3K-Akt pathway to be altered (KEGG Pathway Map, **Supplementary Fig. 7**). The Wnt signaling pathway was also enriched (**Fig. 7B**). Enrichment in the PI3K-Akt and Wnt signaling pathways was of interest given reports that altering membrane cholesterol might affect lipid raft formation and subsequent PI3K signaling (42–44), or that membrane-bound cholesterol can directly activate certain downstream signaling proteins such as Akt or Dvl2 (Wnt signaling).

At the DNA-level, ATAC-seq captured patterns of significantly increased or decreased chromatin accessibility at 1842 and 128 genomic intervals, respectively (**Supplementary Fig. 8**). We show that these dynamic regulatory signatures are proximally enriched close to genes that are downregulated by ABCA1 depletion, indicating a functional link between changes in chromatin state and RNA abundance (**Supplementary Fig. 8**). Integration of differential ATAC and RNA-seq results further captures a significant overrepresentation of dynamic chromatin signatures proximal to genes associated with specific functional terms, such as cholesterol efflux and other cholesterol-related processes (**Supplementary Figure 9**). This would be expected since loss of ABCA1 would lead to a buildup of cholesterol within the cell, engaging cholesterol-related feedback mechanisms (45). In addition, sites with increased chromatin accessibility – enriched for NFκB binding motif signatures – were also significantly overrepresented near genes encoding factors involved in T cell related GO terms, such as negative regulation of T cell co-stimulation (**Supplementary Figs. 8-9**). A neural network based sequence-to-differential-expression model called SEAMoD (46) was trained on the RNAseq and ATACsec data from the macrophages and co-cultured T cells. The model identified multiple transcription factors including Mef2A, Mef2C and GATAs 1-3 as potential drivers of the gene expression differences (**Supplementary Fig. 10**). Many of these transcription factors are also downstream targets of the PI3K/AKT pathway.

A phospho-protein array with 1268 antibodies found changes in the phosphorylation of several proteins, including many implicated in PI3K-AKT signaling, membrane initiated signaling and immune function (**Fig. 7C**). For example, with respect to the PI3K-AKT pathway, we observed phospho-AKT1S1 and TSC2, but a decrease in AKT2. In agreement with our ATAC-seq and RNA-seq analyses, phospho-MEF2A was higher when ABCA1 was knocked down. However, phospho-NFκB (pS518) and phospho-NFκB-p65 (PA311) were slightly decreased.

### AKT subtypes differentially mediate downstream effects of ABCA1 on macrophage function

Since PI3K-AKT signaling was implicated, we evaluated the effect of a PI3K inhibitor (alpelisib) or a pan-AKT inhibitor (capivasertib). As expected, siRNA against ABCA1 in BMDMs reduced subsequent CD8+ T cell expansion (**Fig. 7D**). However, co-treatment of BMDMs with siRNA against ABCA1 and either alpelisib or capivasertib attenuated this effect (**Fig. 7D**). Similarly, either alpelisib or capivasertib were able to rescue the suppressive capacity of BMDMs lacking ABCA1 on cytotoxic T cell activity (**Fig. 7E**). Thus, in terms of BMDMs supporting CD8+ T cell expansion and function, the PI3K-AKT signaling axis is required for, if not mediating, the effects of losing ABCA1.

AKT has three subtypes. To further explore AKT signaling, we made use of selective inhibitors of AKT1 or AKT2 (A-674563 or CCT128930 respectively). Somewhat surprisingly, neither inhibitor was able to attenuate the impaired cytotoxicity of T cells co-cultured with BMDMs lacking ABCA1 (inhibitors co-treated with siRNA against ABCA1, **Fig. 7F**). This indicated that AKT3 might be mediating the downstream effects of ABCA1. In support of this notion, AKT3 protein was increased in BMDMs from mice lacking myeloid expression of ABCA1 (ABCA1^fl/fl^;LysMCre^+^, **Supplementary Fig. 11**).

Less is known about the effects of AKT3 on MΦ function, although it has been implicated in atherosclerosis (47, 48) and tissue remodeling (49). There are no AKT3-selective small molecule inhibitors available. Therefore, to specifically test the involvement of AKT3, we generated BMDMs from control mice (ABCA1^+/+^;LysMCre^+^) or mice lacking ABCA1 expression within MCs (ABCA1^fl/fl^;LysMCre^+^). siRNA against AKT1 in ABCA1^fl/fl^;LysMCre^+^ BMDMs did not significantly alter T cell expansion (**Fig. 7G**). siRNA against AKT2 in ABCA1^fl/fl^;LysMCre^+^ BMDMs did slightly attenuate the effects of losing ABCA1, but not significantly so (**Fig. 7H**). On the other hand, siRNA against AKT3 was able to completely rescue the effects of ABCA1 knockout in BMDMs, in terms of subsequent CD8+ T cell expansion (**Fig. 7I**). Thus, the anti-expansion effects of siRNA against ABCA1 in macrophages on CD8+ T cells requires the activity of PI3K and the presence of AKT3.

On the other hand, efferocytosis continued to be increased when macrophages with knockdown of ABCA1 were treated with a pan-PI3K inhibitor (**Fig. 7J**). However, pan-inhibition of AKT significantly attenuated this response (**Fig. 7J**). Inhibitors to either AKT1 or 2 were able to inhibit the effects of siRNA against ABCA1 on efferocytosis, suggesting that both subtypes were involved (**Fig. 7K**); in stark contrast to the effects of ABCA1 on T cell expansion. Thus, while AKT is a common feature governing the effects of T cell expansion/function and efferocytosis, the specific subtypes differ, as does the dependence on the upstream PI3K.

### Loss of MC-ABCA1 results in increased growth of tumor and metastatic lesions in murine models

In order to determine the combined effects of macrophage-ABCA1, we examined different *in vivo* tumor models. First, 4T1 murine cells were grafted orthotopically and allowed to form tumors in wildtype mice. BMDMs harvested from naïve donor mice were transfected with control siRNA or siRNA against ABCA1. The transfected BMDMs were then grafted into 4T1 tumor bearing mice (experimental setup summarized in **Fig. 8A**). Somewhat surprisingly, given the already aggressive nature of 4T1 tumors, grafting of donor BMDMs with ABCA1 knocked down resulted in significantly increased tumor growth (**Fig. 8B**). As expected, based on our *in vitro* assays, there were fewer CD8+ T cells in tumors grown in mice receiving BMDMs lacking ABCA1 (**Fig 8C**).

**Figure 8:**
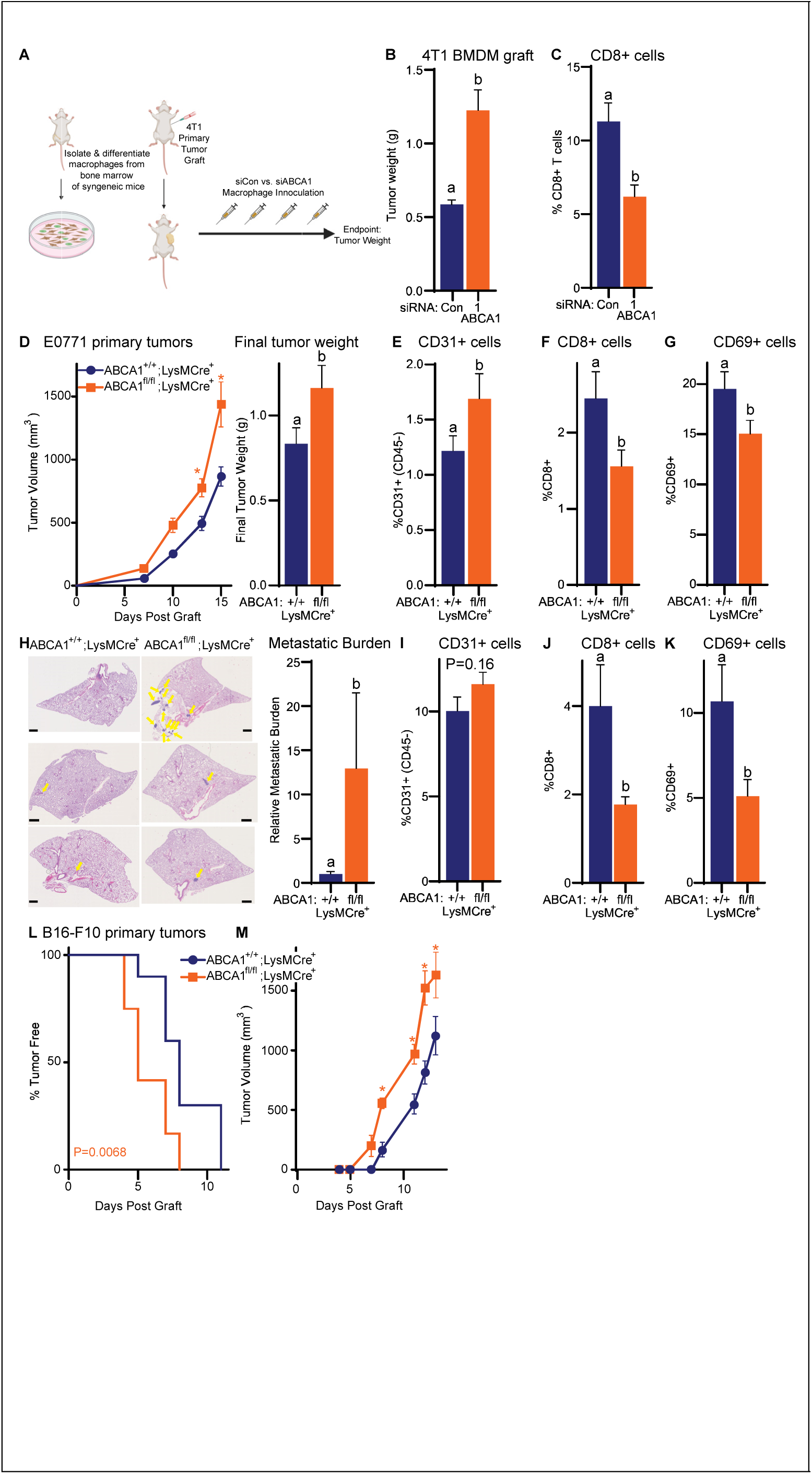
Myeloid cell loss of ABCA1 results in increased tumor and metastatic outgrowth in syngeneic murine models. **(A)** “Cell therapy” with BMDMs transfected with either control or siRNA against ABCA1 were grafted into mice bearing 4T1 mammary tumors. Experimental design is shown (generated and adapted from BioRender). **(B)** Resulting tumor weights are shown (P<0.05, Student’s T Test). **(C)** CD8+ T cells were less abundant in tumors from mice grafted with siABCA1 BMDMs, as assessed by flow cytometry. **(D)** E0771 primary, orthotopic, mammary tumors grew at an increased rate in mice lacking MC-expression of ABCA1 compared to ABCA1-replete mice (ABCA1fl/fl;LysMCre+ vs. ABCA1+/+;LysMCre+ mice respectively). Tumor volumes as measured by calipers through time are presented to the left of final tumor weights at necropsy. **(E)** CD31+ cells were more abundant in tumors grown in ABCA1fl/fl;LysMCre+ mice, while decreases were observed in **(F)** CD8+ T cells and **(G)** cells positive for CD69, a marker of T cell activation CD69. **(H)** Metastatic burden in the lungs was increased in ABCA1fl/fl;LysMCre+ mice 14 days after intravenous graft of E0771 cells. Representative micrographs presented on the left. Scale bar is 800μm. Yellow arrows depict metastatic lesions. Mean total lesion volumes (metastatic burden) quantified and shown on the right. **(I)** CD31+ cells were not significantly increased in tumors lacking MC expression of ABCA1, but **(J)** CD8+ T cells and **(K)** CD69+ cells were decreased. **(L)** Time to detection of a tumor by palpation was significantly decreased and **(M)** subsequent tumors grew faster when B16-F10 melanoma cells were grafted in ABCA1fl/fl;LysMCre+ mice compared to control ABCA1+/+;LysMCre+ mice. Asterisks or different letters denote statistically significant differences.

To more specifically test the role of myeloid cell-ABCA1 on tumor progression, we grafted E0771 murine mammary cells into control mice or those lacking ABCA1 expression specifically in myeloid cells. Tumors grew significantly faster in ABCA1^fl/fl^;LysMcre^+^ mice compared to control (**Fig. 8D**). As expected from our coculture experiments, resulting tumors had increased blood vessel endothelial cells, as denoted by CD31+ staining (**Fig. 8E**). There were also fewer CD8+ T cells (**Fig. 8F**) and decreased CD69+ cells, a marker of immune cell activation (**Fig. 8G**).

Given that metastatic disease is responsible for the majority of breast cancer mortality, we also evaluated this stage of disease by grafting E0771 cells intravenously, effectively modeling colonization and subsequent outgrowth of lesions. Of the mice with noted metastatic disease, the total metastatic burden was significantly higher in ABCA1^fl/fl^;LysMcre^+^ mice compared to control ABCA1^+/+^;LysMcre^+^ mice (**Fig. 8H**). Flow cytometry found increased presence of CD31+ endothelial cells, and decreased CD8+ and CD69+ cells (**Fig. 8I-K**).

Since the effects of ABCA1 within myeloid cells would be expected to be conserved regardless of tumor type, we also evaluated the growth of B16-F0 melanoma tumors. The time to detection of the first palpable tumor was significantly reduced in mice lacking myeloid cell ABCA1 (**Fig. 8L**). Correspondingly, tumor growth through time was increased (**Fig. 8M**).

## DISCUSSION

Tumor associated macrophages have been implicated in several aspects of tumor pathophysiology, including secretion of growth factors, promotion of neoangiogenesis, generalized immune suppression and suppressing T cell function. On the other hand, macrophages are also capable of anti-cancer activities, such as phagocytosis and direct killing of tumor cells, antigen presentation and support of T cell function. We found that ABCA1 activity in macrophages (1) drives them into being better able to infiltrate tumor spheroids, (2) inhibits angiogenesis, and (3) inhibits immune-suppressive efferocytosis without influencing phagocytosis. Furthermore, ABCA1 within macrophages improved their ability to support (a) T cell migration, (b) CD8+ T cell expansion, especially at later divisions, and (c) the ability of CD8+ T cells to kill cancer cells. The cholesterol efflux proteins ABCA1 and ABCG1 have been most investigated in so called ‘foam cells’, cholesterol laden macrophages found in atherosclerotic plaques (50–53), or in the context of retinal diseases such as macular degeneration (54). Many previous studies investigate models of double knockout for ABCA1 and ABCG1, assuming that two efflux proteins operate in a redundant fashion, at least in terms of cholesterol efflux. However, those studies that do knockout them individually suggest that they are not redundant for many, if not most, downstream cellular functions. To date, comprehensive assessment of whether ABCA1 influences different myeloid cell functions has not been performed, especially in the context of tumor biology. Thus, our novel findings open new avenues of research to enhance ICB therapy in breast and other cancers

### Migration/infiltration

Previous work found that double knockout of ABCA1 and ABCG1 in macrophages resulted in decreased migratory capacity when stimulated with C5a (compliment component 5a), compared to wildtype macrophages, ascribed to Rac1 activation (55). We demonstrate that ABCA1 greatly increases macrophage capacity to infiltrate tumor spheroids. This is an important finding as cell-based therapies often require the ability of immune cells to infiltrate tumors or metastatic lesions; in this case, the other beneficial effects of ABCA1 such as T cell support require close contact of macrophages, T cells and the tumor microenvironment (**Fig. 2&5**).

### Angiogenesis

Previous work has demonstrated that LXR induction of ABCA1 reduces angiogenesis, presumably by decreasing lipid raft mediated stabilization of the VEGF-VEGFR complex, within vascular endothelial cells (56). Similarly, angiogenesis is increased in zebrafish embryos lacking ABCA1 and ABCG1 (57). However, endothelial cell knockout of ABCA1 alone did not result in alterations in new vessel sprouting in a murine model of atherosclerosis, while knockout of ABCG1 did (58). Similar to our findings, HMVECs cocultured with BMDMs from myeloid cell conditional ABCA1 knockout mice had increased proliferation (59). Angiogenesis is associated with the classical M2 polarization state of macrophages (60). However, although several aspects of angiogenesis are upregulated in macrophages transfected with siRNA against ABCA1, they are not transitioned to a clear M2-state, at least as determined by gene expression (**Fig. 7**). This reinforces our model whereby ABCA1 shifts macrophages into an alternate polarity, subtly enhancing several aspects of their anti-tumor capabilities. In strong support of our preclinical data, we found three important correlations (**Fig. 3C-E**). First *ABCA1* in macrophages derived from human PBMCs was inversely correlated to their capacity to support tube formation. Second, *ABCA1* was inversely correlated to *VEGFA* in macrophages derived from human macrophages. Finally, when assessing expression from bulk RNA-seq of human breast tumors, an inverse correlation between *ABCA1* and *VEGFA* was observed. Thus, ABCA1 in myeloid cells activity likely represents a clinically important modulator of tumor neo-angiogenesis.

### Efferocytosis

Efferocytosis is a non-immunogenic program mostly used by phagocytic cells engulfing apoptotic bodies. It is seen as a major hurdle to using antigen presenting cells in cell-based therapies (61–64). Interestingly, apoptotic cells increase ABCA1, which is required for the secretion of ‘find me’ signals (65, 66). Within macrophages, ABCA1 seems to protect from the cellular stress associated with engulfing apoptotic bodies, although ABCG1 appeared to be more predominant in this role (67). We have found that ABCA1 activity decreases efferocytosis (**Fig. 4**). The effects were more pronounced when using apoptotic neutrophils as bait compared to cancer cells, perhaps because cancer cells are known to stimulate efferocytotic pathways (68). Significant efforts have been made in trying to identify therapeutic targets to suppress efferocytosis. While some progress has been made with respect to the Fractalkine receptor (CX3C motif chemokine receptor 1), Axl, MerTK, CD47 and Tim4/3, no approaches have thus far been clinically approved (64). Thus, our data reveal ABCA1 as a potential target to curtail efferocytosis in the treatment of solid tumors.

### Support of T cell function

Cholesterol is required to support the rapid clonal expansion of T cells post-activation (39, 40); plasma membrane cholesterol abundance being required for T-cell receptor clustering and signaling (69). Cholesterol homeostasis has also been implicated in tissue resident memory CD8 T cells (70). T cell specific knockout of ABCA1 and ABCG1 mice had decreased T cell numbers but increased activation, resulting in decreased atherosclerosis and aortic inflammation (71). Very little is known about how ABCA1 within myeloid immune cells might influence T cell biology. Using a somewhat contrived system, dendritic cells treated with a statin (zoledronic acid) increased ABCA1 which enhanced extracellular release of the phospho-antigen isopentenyl pyrophosphate, ultimately activating Vγ9Vδ2 T cells (72). Shifting membrane fluidity and lipid rafts could alter the immunological synapse between myeloid cells and T cells or efficiency of membrane-initiated signaling such as through the toll-like receptors (73). Indeed, we find evidence of more open chromatin around predicted NFκB binding sites when ABCA1 is knocked down (**Fig. 7**, **Supplementary Fig. 8**). However, antigen presentation was not required since CD3/28 activated T cells also showed altered expansion when ABCA1 was altered in macrophages. On the other hand, direct contact between macrophages and T cells was required for the full effects observed **(Fig. 5F, Supplementary Fig. 4I).**

Dendritic cells lacking both ABCA1 and ABCG1 have enhanced inflammasome activation, ultimately resulting in enhanced T cell activation and polarization to the T_h_1 and T_h_17 types (74). These findings from double knockout of ABCA1 and ABCG1 are contrary to our findings in macrophages where only ABCA1 is knocked down or inhibited (**Fig. 5**). We find that loss of ABCA1 in macrophages impairs CD8+ T cell expansion while overexpression enhances it. These robust effects appear to require direct contact between macrophages and T cells, although some enhancement was observed when the cell types were isolated by a membrane (**Fig. 5**, **Supplementary Fig. 4**). Furthermore, the resulting expanded CD8+ T cells had less or more anti-cancer cytotoxic capacity when cultured with macrophages lacking or overexpressing ABCA1 respectively. These data provide more support that ABCA1 and ABCG1 are not redundant, and may even have opposing effects. It is also possible that myeloid cell type or context may dictate the effects of ABCA1.

In human breast tumors, *ABCA1* expression was positively correlated with CD8 T cell abundance as well as enzymes indicative of CD8 T cell activity (*IFNγ*, *GZMB* and *PRF1*; **Fig. 6**). mIF analysis of breast tumor microarrays also demonstrated a correlation between ABCA1 and CD8 staining intensity. Importantly, this correlation held in the relationship between myeloid cell expression of ABCA1 and the number of CD8+ cells. Thus, our data indicating that myeloid cell ABCA1 activity supports CD8 T cell expansion and function are likely clinically relevant, and position ABCA1 as a potential therapeutic target.

### Myeloid cell ABCA1 and tumor growth

Together, our data strongly suggest that ABCA1 in murine macrophages skews them towards a robust anti-cancer phenotype. Syngeneic mammary and melanoma tumors grew at a significantly quicker rate in mice with conditional knockout of *Abca1* in myeloid cells (**Fig. 8**). Likewise, 4T1 mammary tumors grew to a larger size in mice receiving supplementation with adoptive macrophages transfected with siRNA against *Abca1*. These data are strongly supported by correlational data in humans, demonstrating that elevated ABCA1 is associated with (1) improved response to ICB, (2) decreased *VEGFA* angiogenesis, and (3) increased CD8+ cell expansion and infiltration in tumors. However, a previous report indicated that myeloid cell loss of ABCA1 decreased B16-F10 melanoma and MB49 bladder tumor growth (75). Knockdown of the related transporter ABCG1 also resulted in inhibited tumor growth (75), but in a different study this required a physiologic challenge (western diet, age or genetic model of hypercholesterolemia) (76). Another report suggested that ovarian tumor cells secrete some factor that upregulates ABCA1 and ABCG1 in macrophages, which then polarized them into a tumor-associated or “M2-like” state (77). It is not clear why there is discrepancy between reports. Variables to consider include genetic drift of the knockout mouse line and cancer cells, age of mice, and potentially other environmental factors (like season, stress etc).

### ABCA1-influenced functions utilize different aspects of the PI3K-AKT signaling pathway

Cholesterol within the plasma membrane helps govern membrane fluidity. It is also a key component of lipid rafts; lipid rafts being able to bring membrane associated proteins within close proximity to each other, enhancing signaling. One example of this comes from tumors grown in hypercholesterolemic mice (ApoE^-/-^), which have increased PI3K and AKT signaling, and treatment with a PI3K inhibitor was able to attenuate growth (78). Another example comes from colorectal cancer where the APC oncogene increased plasma membrane cholesterol and thus WNT signaling hubs (lipid rafts) (38).

We found that loss of ABCA1 in macrophages impairs subsequent CD8+ T cell function in a PI3K dependent fashion (**Fig. 7**). Thus, we propose a model whereby loss of ABCA1 results in increased membrane cholesterol and hence lipid rafts and associated PI3K activity. Interestingly, AKT1 was not found to mediate the effects of PI3K with respect to subsequent T cell support, despite being previously linked to polarizing macrophages towards an M2 state. Specifically, PI3K-AKT dependent MerTK activity inhibited polarization towards an M1 state (79). Instead, our data suggest that AKT3 was involved. This is the first description of AKT3 activity in macrophages resulting in altered support of lymphocyte activity.

Given the known immune-suppressive pathways engaged through efferocytosis, we speculated that the observed regulation efferocytosis by ABCA1 activity may be related to the effects of ABCA1 expression on T cell expansion and function. Indeed, previous reports have indicated that PI3K modulates efferocytosis (80) However, a PI3K inhibitor did not attenuate the effects of siABCA1 on efferocytosis, whereas PI3K inhibition did reduce the effects of siABCA1 on T cell expansion and cytotoxic functions. The independence of efferocytosis from PI3K is similar to a previous report where efferocytosis was not affected by treatment of alveolar macrophages from patients with chronic obstructive pulmonary disease (COPD) with PI3K inhibitors (81). We found that AKT1 and 2 are likely involved in mediating the effects of losing ABCA1 on efferocytosis by macrophages. Recent reports suggest that certain PZD-domain containing proteins such as Dvl2 can be directly activated by cholesterol within the inner leaflet of the plasma membrane (43, 44, 82–84). Thus, one mechanism may be accumulation of cholesterol within the inner leaflet of the plasma membrane when ABCA1 is absent/compromised (**Supplementary Fig. 1H**), facilitating direct interactions with AKT1 and 2. Other putative mechanisms include lipid raft formation and promotion of other signaling pathways that cross-talk with AKT1/2.

### Conclusions

We demonstrate that the activity of the cholesterol efflux and translocation protein, ABCA1, has critical implications in the regulation of various myeloid cell functions, including tumor infiltration, efferocytosis, and the support of T cell expansion and anti-cancer cytotoxic activities. Correlational data from human breast tumors strongly support our preclinical observations. These data provide further mechanistic insight into how cholesterol homeostasis influences the progression of breast and other tumors. Tumors grown in mice lacking ABCA1 in myeloid cells grew at an accelerated rate, likely the culmination of the many different myeloid cell functions that ABCA1 activity modulates. Thus, by regulating several different anti-tumor associated functions in myeloid cells, ABCA1 represents an upstream node, ripe for therapeutic exploitation.

## Materials and Methods

### Reagents

PSC833 was purchased from Sigma Aldrich (Cat No 121584-18-7) and dissolved in DMSO. OVA peptide (257-264, chicken) was purchased from Sigma Aldrich (Cat No S7951-1MG) and dissolved in PBS, anti-CD3 and anti-CD28 antibodies were purchased from BD Biosciences, Cat No 553057 and 553294 respectively. LPS was purchased from ThermoFisher (Cat No: 00-4973-03). Antibodies for flow cytometry were purchased from BD Biosciences and ThermoFisher. FACS Buffer is 2% Fetal Bovine Serum. CFSE was purchased from Biolegend, Cell Trace Red, and Cell Trace Violet from ThermoFisher. FBS was from Gibco Cytvia HyClone. Non-essential amino acids, Sodium Pyruvate, Penicillin/Streptomycin, PBS, cell stripper solution, RPMI-1640 and trypsin were from Corning. DMEM was from Fisher Sciences. siRNAs against ABCA1 (siRNA ID: SASI_Mm01_00075553, SASI_Mm01_00075555), AKT1(siRNA ID: SASI_Mm02_00316647), AKT2 (siRNA ID: SASI_Mm02_00304936) and AKT3(siRNA ID: SASI_Mm01_00200790) were purchased from Sigma Aldrich.

### Cell Culture

E0771 and 4T1-luc cells were a gift from Mark Dewhirst (Duke University) and cultured in DMEM F12 (ThermoFisher) supplemented with 10% FBS, 1% non-essential amino acids, 1% sodium pyruvate and 1% penicillin/streptomycin. E0771-OVA cells were a gift from Jun Yan (University of Louisville) and grown in the above-described conditions. All cell lines were tested for Mycoplasma contamination.

### Isolation of Bone Marrow-Derived Macrophages (BMDMs)

Bone marrow cells were collected from the mouse tibia and femur by flushing with complete DMEM. Cells were then passed through a 70μm filter. Complete Macrophage Media consisting of DMEM supplemented with 25 ng/mL of M-CSF (PeproTech) was added on day 2, and media was replaced on day 6 and every 2-3 days after that. BMDMs were harvested between day 10 – day 12 using cell stripper solution and were used for subsequent experiments.

### Animal Studies

All animal protocols were approved by the Institutional Animal Care and Use Committee (IACUC) at the University of Illinois at Urbana-Champaign. BALB/C and C57BL6/J mice were purchased from either Charles River Laboratories or The Jackson Laboratory. ABCA1^fl/fl^ and LysMCre+ mice were purchased from The Jackson Laboratory, and bred at the University of Illinois to obtain ABCA1^fl/fl^;LysMCre^+^ and ABCA1^+/+^;LysMCre^+^ mice. Mice were housed with ad libitum access to food and water and 12 h light and dark cycles.

### In-Vivo Macrophage Infiltration Study

300,000 4T1 cells were grafted orthotopically in the mammary fat pad and were allowed to grow for 3 days. BMDMs harvested from syngeneic mice were transfected with either a control or siRNA against ABCA1 which were then injected in the subcutaneous space for consecutive 4 days. Tumors were then weighed post-euthanasia and analyzed via flow cytometry.

### ABCA1^fl/fl^;LysMCre^+^ E0771 Primary Tumor Study

We obtained ABCA1*^fl/fl^* mice from The Jackson Laboratory and bred them with LysMCre + mice to generate a myeloid-specific ABCA1 knockout cell line. These mice were genotyped using the protocol provided by the Jackson Laboratory. 200,000 E0771 cells were grafted in the mammary fat pad of age-matched females 2-5 months old. Tumors were measured over time using a digital caliper. Tumors were then weighed post-euthanasia and analyzed via flow cytometry.

### ABCA1^fl/fl^;LysMCre^+^ E0771 Metastasis Study

150,000 E0771 cells were injected intravenously via the tail vein into age-matched females 2-5 months old. 2 weeks post graft, mice were euthanized. Lungs were analyzed using histology and flow cytometry.

### ABCA1 ^fl/fl^;LysMCre^+^ B16-F10 Primary Tumor Study

200,000 B16-F10 cells were grafted in the mammary fat pad of age-matched females 2-5 months old. Tumors were measured over time using a digital caliper. Tumors were then weighed post-euthanasia and analyzed via flow cytometry.

### Gene Silencing and Overexpression

GenMute transfection reagent (SignaGen) for primary macrophages was used for siRNA-mediated knockdowns. Effectene transfection reagent (Qiagen) was used for over-expression. Media was replaced 4 hours after transfection and all downstream analysis was carried out 48 hours after transfection.

### Flow cytometry staining and analysis

For tissue samples, fresh tissues were collected from mice and digested in DMEM F12 supplemented by 2 mg/ml type II collagenase and 1 % penicillin/streptomycin for 30-60 min at 37°C while shaking. Cells were then passed through a 70μm filter, washed with FACS, incubated with ACK lysis buffer followed by another wash with FACS Buffer. For surface staining, cells were fluorescently conjugated antibodies in FACS (PBS supplemented with 2% FBS and 1% penicillin-streptomycin) at a concentration of 1:50-1:100 for 30 – 60 minutes, followed by FACS washes and fixed in 4% formaldehyde if needed. Intracellular staining was carried out using BD Fix/perm reagents according to the manufacturer’s instructions. FCS Express and companion software for the Thermo Attune instrument were used to capture and analyze data from flow cytometry.

### T cell Migration Assay

A pre-optimized amount of BMDMs were seeded and primed with OVA and LPS. Pan T Cells were isolated from the spleen of OT-1 mice and co-cultured with pre-primed and pre-treated BMDMs for 72 hours. E0771-OVA cells were seeded on a 24-well plate. Post co-culture with BMDMs, T Cells were stained with cFSE, and a pre-determined amount of T Cells were added in the top chamber of a Boyden chamber with 3.0 µ pore size, T Cells were allowed to migrate for 10 hours. Cells were then lysed using Triton-X, fluorescence of migrated cells was measured using plate reader.

### T Cell Expansion Assay

A pre-optimized number of BMDMs was seeded followed by treatment or transfection. Pan T Cells were isolated from the spleen of wildtype C57BL6 mice using Untouched mouse T Cells Kit (Dynabeads), stained with cFSE (biolegend), and activated using antiCD3, antiCD28 antibodies (Biolegend). T Cells were co-cultured with pre-treated BMDMs for 72 hours, followed by staining of surface protein markers. Samples were analyzed using flow cytometry.

### T Cell Cytolysis Assay

A pre-optimized number of BMDMs was seeded and primed with OVA peptide and LPS. Pan T Cells were isolated from the spleen of OT-1 mice and co-cultured with pre-primed BMDMs for 72 hours. E0771 and E0771-OVA cells were labeled with cFSE and Cell Trace Far Red or Cell Trace Violet respectively and then plated in a 24-well plate. Subsequently, activated T Cells were counted and co-cultured with cancer cells at various ratios. 24 hours after co-culture, cytolytic activity was assessed by flow cytometry. The ratio between OVA-expressing cells and Parental cells was determined.

### Organoid Infiltration Assay

1.5% agarose was cast in 96 well u bottom ultra-low attachment plates. 4T1 or E0771 cells were pre-stained with HCS Nuclear Mask Blue (ThermoFIsher, Cat No: H10325) and Cell Mask Orange (Cat No: C10045), and layered on top of agarose gel in each well. Tumorspheres were cultured for 7 days, media was replaced every 3 days. On day 7, pre-treated BMDMs were stained with Cell Trace Red (ThermoFisher, Cat No: C34572) and resuspended and layered on top of organoids and allowed to infiltrate overnight. Next day, all media in the well was replaced with Live Cell Imaging solution (ThermoFisher, Cat No: A59688DJ), and tumor spheres were imaged using LSM900 Confocal Microscope.

### HUVEC Tube Formation Assay

HUVECs were procured from ATCC and cultured in Human Large Vessel Endothelial Cell Basal Medium (ThermoFisher) supplemented with Large Vessel Endothelial Supplement (LEVS) as per the manufacturer’s instructions and 1% penicillin-streptomycin. 100uL Geltrex (ThermoFisher) was evenly applied on a 24-well glass bottom plate and HUVEC cells were layered on top. After 4 hours, pre-treated BMDMs were co-cultured with HUVECs using a 0.4μm Boyden chamber overnight. Post-co-culture HUVECs were stained with Calcein AM dye (ThermoFisher) and imaged using an LSM900 confocal microscope.

### Efferocytosis Assay

BMDMs were stained with Cell Tracer Red (ThermoFisher), treated, and seeded, at 80% confluency. Polymorphonuclear Neutrophils (PMNs) were isolated from the bone marrow of wildtype mice using Ly6G isolation beads (MiltenyiBiotec) as per manufacturer’s instructions, stained with CFSE (Biolegend), and treated with staurosporine (Sigma Aldrich) for 4-6 hrs. Post-treatment, PMNs were washed with PBS twice and co-cultured overnight with BMDMs at a 1:2 BMDM:PMN ratio. Cells were analyzed using flow cytometry.

### Phagocytosis Assay

A pre-determined amount of BMDMs were seeded, and transfected or treated as needed. Phagocytosis Assay Kit was purchased from Cayman Chemicals and the experiment was carried out as per the manufacturer’s instructions. Data was analyzed using Flow Cytometry.

### Human PBMC Analysis

Blood Samples from female volunteers was obtained, mononuclear cells were obtained by density gradient centrifugation using Lymphoprep medium (StemCell Cat No: 18060). T Cells were isolated using Pan T Cell isolation kit from Miltenyi as per manufacturer’s instructions. T Cells once isolated were stored in Cytostor medium. Flowthrough from T Cell isolation was cultured in X-Vivo Medium supplemented with human m-csf (Peprotech) for 7 days to obtain mature human macrophages (hMDMs). T Cells were stained with cFSE and co-cultured with patient matched hMDMs for 72 hours followed by surface staining and flow cytometric analysis.

### RNA Sequencing and Data Analysis

Bulk RNA sequencing was conducted on an SP lane with 1×100nt reads, generating more than 480 million reads. DEG list was used to plot the heatmap and perform Gene Set Enrichment Analysis and Gene Ontology Analysis.

### ATAC Sequencing and Data Analysis

ATAC-seq fastq files were trimmed using Trim Galore (paired-end, quality score ≥ 30). Trimmed reads were aligned to the mouse genome (GRCm39) using Bowtie2, and the sorted and indexed BAM files created using Samtools. Peak calling was performed with MACS2 using the parameters --bdg --nomodel --g mm to identify accessible chromatin regions in each sample. To combine peaks across all samples, Bedtools merge was used to create a merged peak set, and coverage signal across all merged peaks was quantified with Bedtools coverage. Read counts were normalized to reads per million mapped reads (RPM), with the total number of mapped reads calculated using Samtools view (-c -F 256). Differential accessibility analysis was carried out using DESeq2 on three biological replicates of macrophage control and treatment samples, to identify differentially accessible regions. For motif enrichment analysis, peaks were divided into upregulated and downregulated groups based on the SWFC (significance weight fold change) score (-log10(p_adj) * log2(Fold Change)), with thresholds of -1 and 1 for upregulated and downregulated peaks, respectively. Genomic sequences of these peak groups were extracted using Bedtools getfasta, and motif enrichment was assessed using SEA (-- thresh 10.0 - -align center) from the MEME Suite with the HOCOMOCOv11_full_mouse_mono motif database. Non-differential peaks (SFW between -1 and 1) were used as a control for motif enrichment analysis.

For differential ATAC-seq peak enrichment relative to gene expression signatures, transcript coordinates were obtained from BioMart. Genes were classified into upregulated, non-regulated, and downregulated categories based on their SWFC scores, using a threshold of ±0.8. Gene Ontology (GO) analysis was performed using GeneOntology.org to identify functional enrichments in these gene sets. To assess the enrichment of ATAC-seq peaks near upregulated and downregulated genes, the density of ATAC peaks was calculated for each gene set (upregulated and downregulated) across various window sizes ranging from 10 kb to 50 Mb, using the formula: density = (10^8^) * number of ATAC peaks / window size. The expected density for each set was calculated by permutation (x1000) against an equal-sized random gene set. This analysis was performed separately for both the upregulated and the downregulated ATAC peaks.

For differential expression enrichment of Gene Ontology (GO) terms, the mean SWFC score of genes within each GO term was calculated. The Z-score of enrichment for each GO term was determined by comparing the observed mean SWFC score to a null distribution generated through 1000 random permutations of similarly sized gene sets. To assess the nearby enrichment of upregulated ATAC-seq peaks in relation to GO terms, only the downregulated GO terms (SWFC < (-1)) were selected, as the downregulated genes showed closer proximity to the upregulated ATAC peaks in the above method. The density of ATAC peaks(upregulated) was calculated around the gene sets for each selected GO term using the formula: Density=(10^6^)×number of ATAC peaks/window size. The Z-score for each GO term was then calculated by comparing the observed ATAC peak density to a null distribution, which was created through 1000 random permutations of equal-sized random gene sets.

We used a variant of the SEAMoD tool for motif discovery (46). This tool uses an interpretable neural network model to relate cis-regulatory sequences (promoters and enhancers) to gene expression in varying cellular contexts. Motifs are learnt as convolutional filters that are parameters of the model. SEAMoD protocol was slightly modified to accommodate the data sparsity. Genes and the set of differentially expressed genes (DEGs) were split into three sets - training, validation and test. A DEG was assigned a value of 1 or -1 to indicate up and down regulation where as 0 if the expression didn’t show a significant change in expression (these values make the model output). For each gene the set 4 nearest ATAC peaks (within 100kbp from the transcription start site) in a cell type-condition pair were selected as the putative cis-regulatory modules (CRMs). All genes from the training set were used to train the model in the first round. Nearest accessible ATAC peak (nearest neighbor) for a gene in each cell type-condition pair were selected as the CRMs for the first round. This was followed by another round of model training using only those genes which were differentially expressed in at least one of the cell types and the CRMs were kept the same as the first round. Next the set of best CRMs (based on the model from the second training round) were identified (46). The model was trained for the third time using all genes from the training set and the set of best enhancers to fix the convolutional filters. Remaining trainable parameters were further fine-tuned by fourth round of model training (from the previous step) using only the DEGs while keeping the convolutional filters unchanged. This final model was then used to do all further analysis. In-silico knockdown was performed (same as from (46)) to obtain the convolutional filters that were most responsible for the model’s predictions. These filters were then converted into position weight matrices (PWMs) using the same protocol as described (46). The PWMs thus obtained were then compared to the a compendium of previously characterized PWMs in mouse using TOMTOM (85) to obtain the final list of important PWMs and associated TFs. Motifs with TOMTOM q-value less than 1 were selected. The PWM logos in **Supplementary Fig. 10** were generated using ggseqlogo (86).

## Supporting information

Bendre et al Supplementary Figures

## Acknowledgments

We would like to thank the patients whose tumors populated data in the TCGA, METABRIC, Human Protein Atlas initiatives, and breast tumor microarrays. We would also like to thank our breast cancer advocate team: Sarah Adams, Renaé Strawbridge, Jamie Holloway, Lea Ann Carson, Susan Stewart and Catherine Applegate. We are grateful for bioinformatic analysis and support from Jenny Drnevich (High-Performance Biological Computing, University of Illinois). The Tumor Engineering and Phenotyping Core at the Cancer Center at Illinois assisted with histology (with thanks to Renee Walker). The Roy J. Carver Biotechnology Center performed the RNA-sequencing.

## Funding

Department of Defense Era of Hope Scholar Award BC200206/W81XWH-20-BCRP-EOHS (ERN)

National Institutes of Health grant R01 CA288207 (ERN)

National Institutes of Health grant R01 CA234025 (ERN)

National Institutes of Health grant T32 GM136629 (HEVG)

National Institutes of Health grant T32 ES007326 (ATN)

National Institutes of Health grant T32 EB019944 (CPS)

The Endocrine Society Research Experiences for Graduate and Medical Students Award (SVB)

The Cancer Center at Illinois Graduate Cancer Scholarship Program Award (SVB)

Postdoctoral Fellows Program at the Beckman Institute for Advanced Science and Technology (NK)

University of Illinois (KVB, ET, ERN)

The State of Texas Grant for Rare and Aggressive Breast Cancer (WAW)

National Institutes of Health grant R01 CA264529 (WAW)

National Institutes of Health grant R01 CA284102 (WAW)

## Supplementary Figure Legends

**Supplementary Figure 1.**

**(A)** Cluster map from single cell RNA-sequencing indicating *ABCA1* expression in healthy, human breast tissue. (data from the Human Protein Atlas; corresponding cluster map in **Fig. 1**). **(B)** *Abca1* expression in different myeloid immune cells isolated from naïve and tumor-bearing (TBM) mice. BMDMs were polarized to M1 with LPS and IFNγ and to M2 by IL4 and IL13 for 48 hours. **(C-D)** Two different siRNAs against ABCA1 result in the expected decrease in ABCA1 mRNA expression in BMDMs. **(E)** Overexpression of ABCA1 resulted in the expected increase in ABCA1 compared to empty vector (pcDNA) control transfected BMDMs. **(F)** ABCA1 protein expression as determined by Western Blot analysis, indicating successful knockdown upon transfection of BMDMs with two different siRNAs against ABCA1 (representative images and quantified band density to the right). **(G)** siRNA against ABCA1 decreased cellular efflux of cholesterol in BMDMs. **(H)** siRNA against ABCA1 also led to an increase in cholesterol within the inner leaflet of the plasma membrane. A ratiometric fluorescent cholesterol probe was injected intracellularly and cells imaged. Representative images to the left of quantified data. [chol]i is the relative concentration of cholesterol within the inner leaflet of the plasma membrane. For C-E, G and H, different letters denote statistically significant differences.

**Supplementary Figure 2: *Knockdown of ABCA1 in macrophages decreases their ability to infiltrate tumor spheroids***. BMDMs were transfected with control or siRNA against ABCA1 for 48h prior to being stained with Cell Trace Far Red and overlayed onto 4T1 tumor spheroids previously stained with PKH26 (Sigma Aldrich), in an ultra-low attachment 96-well plate. Spheroids were then removed, disrupted mechanically and quantified by flow cytometry. Different letters denote a statistically significant difference (P<0.05, Student’s T Test).

**Supplementary Figure 3: *ABCA1 inhibitor PSC833 modestly increases ability of macrophages to support tube formation by HUVEC cells***. Representative micrographs shown. Data is in support of **Fig. 3.**

**Supplementary Figure 4: *ABCA1 activity results in BMDMs that support T cell migration, expansion.*** (A) CD8+ T cell expansion is decreased when activated T cells are co-cultured with BMDMs transfected with siRNA against ABCA1. The percentage in each clonal generation shown here. Data in support of **Fig. 5C**. **(B)** A second siRNA against ABCA1 yielded similar results. **(C)** The effects of siRNA against ABCA1 in BMDMs on T cell expansion were restricted to CD8+ T cells. When CD4+ T cell expansion was assessed, no significant differences were observed (same experiment as in **Fig. 5C** where CD8+ expansion is shown). A ratio between the proportion of CD4+ T cells expanded under treatment conditions to vehicle control conditions is shown to the left, with the blue line being siControl conditions. The corresponding percentage in each clonal generation is shown to the right. **(D)** CD8+ T cell expansion is decreased when activated T cells are co-cultured with BMDMs pretreated with the ABCA1 inhibitor, PSC833. The percentage in each clonal generation is shown. Data in support of **Fig. 5D**. **(E)** Overexpression of ABCA1 in BMDMs enhanced CD8+ T cell expansion. The percentage in each clonal generation shown here. Data in support of **Fig. 5E**. **(F)** BMDMs were transfected with either control or an ABCA1 overexpression vector. 48 hours post-transfection, BMDMs were incubated in 50µM cholesterol for 24 hours prior to being washed and then co-cultured with activated T cells. Left: Resulting CD8+ T cell expansion after BMDMs were treated with cholesterol solubilized in DMSO. The percentage in each clonal generation shown. Right: Resulting CD8+ T cell expansion after BMDMs were treated with cholesterol solubilized in cyclodextrin. The percentage in each clonal generation shown. **(G)** The robust decreases in CD8+ T cell expansion when pan-T cells were cultured with BMDMs lacking ABCA1 is lost when only CD8+ T cells are cultured with BMDMs. CD8+ T cells were bead-isolated prior to incubation with BMDMs that had previously been transfected with control or two different siRNAs against ABCA1. Resulting CD8+ T cell expansion was assessed. The percentage in each clonal generation shown. **(H)** Similarly, the increased CD8+ T cell expansion observed when pan-T cells are co-cultured with BMDMs overexpressing ABCA1 is muted when only CD8+ T cells are cultured with BMDMs. The percentage in each clonal generation is shown. **(I-J)** The robust effects of BMDM-ABCA1 on CD8+ T cell expansion are attenuated when contact between BMDMs and T cells is prevented through the use of a Boyden chamber. The percentage in each clonal generation after BMDMs were transfected with two different siRNAs against ABCA1 is shown in **I**. The percentage in each clonal generation after BMDMs were transfected with an ABCA1 expression vector is shown in **J**.

**Supplementary Figure 5: *ABCA1 staining is associated with CD8 staining in human breast tumor tissue microarrays.*** The data in this supplementary figure are in support of **Fig. 6**. Human breast tumor tissue microarrays (TMAs) were co-stained for ABCA1, the pan-myeloid cell marker CD11B, the cytotoxic T cell marker CD8, and the nuclear stain DAPI, in a mIF assay. Staining intensity as well as % cells positive for each stain was quantified. **(A)** A few cases of normal breast and tumor-adjacent normal breast tissue were included in some of the TMAs. Here, ABCA1 staining intensity is shown when parsed this way. We had too few samples to power the statistical analysis for this. **(B)** There is a positive correlation between ABCA1 and CD8 staining intensity (Spearman coefficient and P values shown). **(C)** There is a positive correlation between percentage of cells co-staining for CD11B and ABCA1, and the percentage of CD8+ cells. Spearman coefficient and P values shown. **(D-E**) Data for B & C were parsed based on ABCA1 into quartiles. Different letters denote statistical difference from one-another (P<0.05; Kruskal-Wallace followed by Dunn’s multiple comparison test). **(F-H)** Samples for each different breast cancer subtypes were assessed independently, evaluating staining intensity. Spearman coefficients and P values shown for F&G while the Pearson coefficient and P value is shown for H. **(I-K**) Data for F-H were parsed based on ABCA1 into quartiles. Different letters denote statistical difference from one-another (P<0.05; Kruskal-Wallace followed by Dunn’s multiple comparison test, with the exception of K, where a 1-Way ANOVA was used with the P value indicated). **(L-N**) Samples for each different breast cancer subtypes were assessed independently, evaluating the percentage of positive cells. Spearman coefficient and P values shown. **(O-Q)** Data for L-N were parsed based on percentage of CD11B+;ABCA1+ cells into quartiles. Different letters denote statistical difference from one-another (P<0.05; Kruskal-Wallace followed by Dunn’s multiple comparison test, with the exception of Q, where a 1-Way ANOVA was used with the P value indicated).

**Supplementary Figure 6: *GO analysis of DEGs between control and siRNA-ABCA1 transfected BMDMs***. Gene Ontology (GO) was performed on the differentially expressed genes (DEGs) identified from RNA-Seq. GO enrichment was calculated and visualized to highlight the most significantly enriched GO Terms using ShinyGO.

**Supplementary Figure 7: *KEGG Pathway analysis of DEGs DEGs between control and siRNA-ABCA1 transfected BMDMs***. Gene Set Enrichment Analysis (GSEA) for KEGG Pathway analysis was conducted on DEGs from RNA-Seq using the clusterProfiler package in R. Results were visualized using GSEA plots and enrichment maps to highlight significantly enriched KEGG pathways.

**Supplementary Figure 8: *ATAC-seq analysis of BMDMs transfected with control or siRNA against ABCA1***. Differential ATAC-seq and enrichment proximal to dynamic gene signatures following ABCA1 depletion. <u>Left</u>: Volcano plot analysis of significantly upregulated (red, 1,842 peaks) and downregulated (blue, 128 peaks) following ABCA1 knockdown. Inset highlights top 3 motifs enriched in upregulated ATAC peaks (SEA analysis; MEME suite). <u>Right</u>: Proximity-based enrichment of upregulated ATAC peaks (top, red) and downregulated peaks (bottom, blue) relative to significant upregulated genes (red) and downregulated genes (blue). Observed densities (solid) are shown in relation to expected densities (dashed).

**Supplementary Figure 9: *ATAC-seq analysis of BMDMs transfected with control or siRNA against ABCA1***. Proximity-based enrichment of upregulated ATAC peaks relative to specific gene sets, defined by shared gene ontology (GO). Z-score ATAC density (y-axis) represents the proximal density of upregulated ATAC peaks (50 Kb, left; 500 Kb, right) compared against a randomized null model. -Z score GO gene set (x-axis) represents the mean significance weighted fold change (SWFC) for a given gene set (with shared GO term) compared against a randomized null model. GO term:: Gene sets related to cholesterol (magenta) and T cell (green) are highlighted, with top GO term enrichments for “Negative Regulation of T cell costimulation” (50 Kb) and “Cholesterol Efflux” (500 Kb) labeled.

**Supplementary Figure 10: SEAMoD *Nueral network learning using RNAseq and ATACseq datasets identified transcription factors likely to drive the changes in differentially expressed genes.*** Motifs obtained from the SEAMoD model trained on the data from the macrophages (M) and co-cultured T cells (T). Convolutional filters were converted into position weight matrices (PWMs), which were then matched with a compendium of previously characterized motifs in mouse using TOMTOM software. Only those motifs were selected that were found significant by the model in one or both cell types and had a strong match with known mouse motifs (see methods). Note that only top 5 Mouse motif matches are shown here. Missing motifs indicate that either no or fewer than 5 matching motifs were found.

**Supplementary Figure 11: *AKT3 protein expression is increased in BMDMs lacking ABCA1***. Representative Western blot AKT3 comparing BMDMs from ABCA1+/+;LysMCre+ mice to BMDMs from ABCA1fl/fl;LysMCre+ mice.

