## Supplementary material for "Cholesterol efflux protein, ABCA1, supports anti-cancer functions of myeloid immune cells": Bendre et al Supplementary Figures

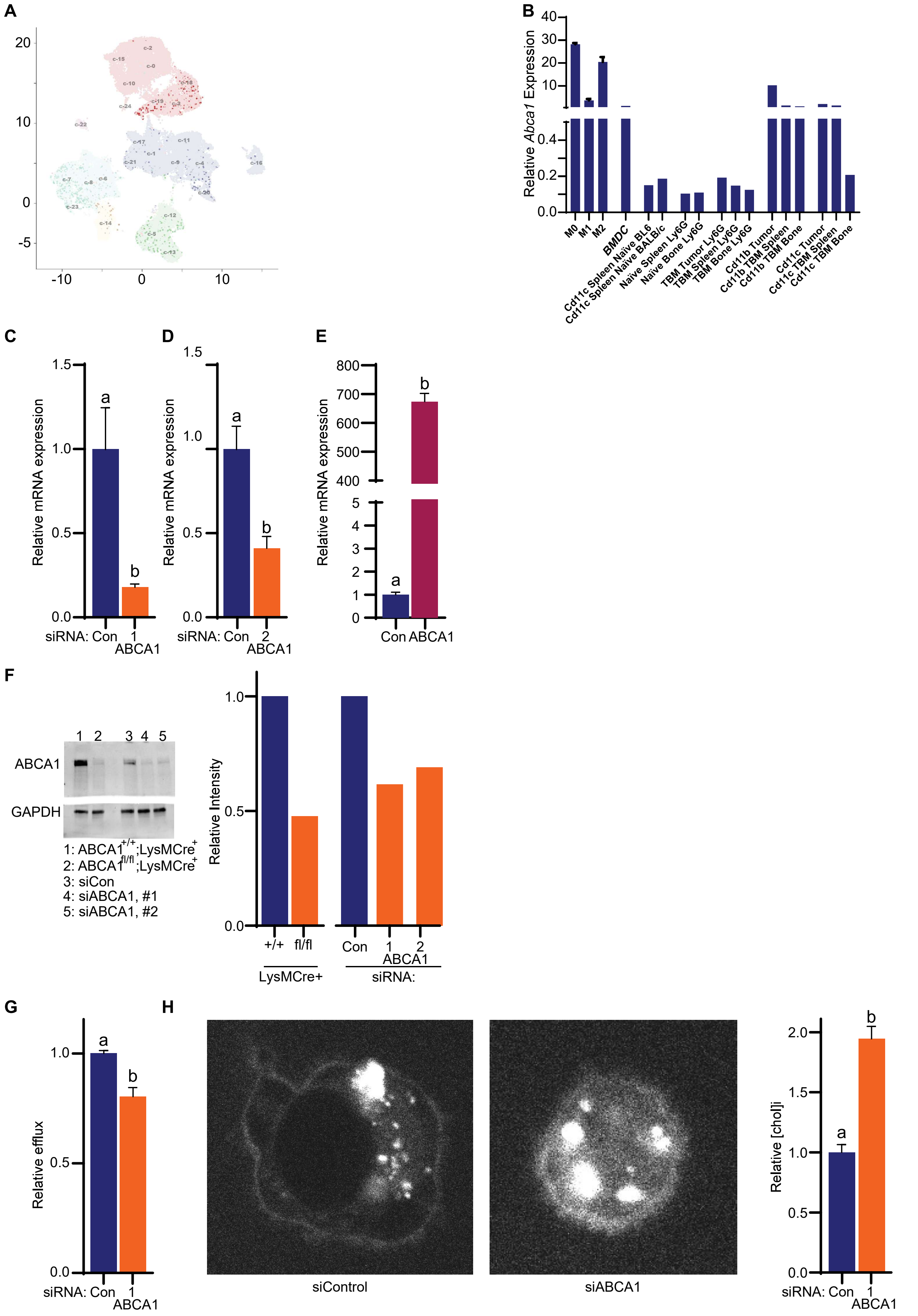


Supplementary Fig. 1

**Supplementary Figure 1.**
**(A)** Cluster map from single cell RNA-sequencing indicating *ABCA1* expression in healthy, human breast tissue. (data from the Human Protein Atlas; corresponding cluster map in **Fig. 1**). **(B)** *Abca1* expression in different myeloid immune cells isolated from naïve and tumor-bearing (TBM) mice. BMDMs were polarized to M1 with LPS and IFN$\gamma$ and to M2 by IL4 and IL13 for 48 hours. **(C-D)** Two different siRNAs against ABCA1 result in the expected decrease in ABCA1 mRNA expression in BMDMs. **(E)** Overexpression of ABCA1 resulted in the expected increase in ABCA1 compared to empty vector (pcDNA) control transfected BMDMs. **(F)** ABCA1 protein expression as determined by Western Blot analysis, indicating successful knockdown upon transfection of BMDMs with two different siRNAs against ABCA1 (representative images and quantified band density to the right). **(G)** siRNA against ABCA1 decreased cellular efflux of cholesterol in BMDMs. **(H)** siRNA against ABCA1 also led to an increase in cholesterol within the inner leaflet of the plasma membrane. A ratiometric fluorescent cholesterol probe was injected intracellularly and cells imaged. Representative images to the left of quantified data. [chol]i is the relative concentration of cholesterol within the inner leaflet of the plasma membrane. For C-E, G and H, different letters denote statistically significant differences.

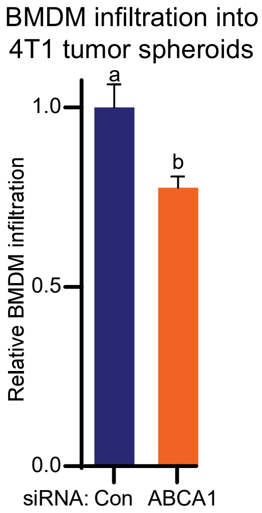


Supplementary Fig. 2

**Supplementary Figure 2:** ***Knockdown of ABCA1 in macrophages decreases their ability to infiltrate tumor spheroids***. BMDMs were transfected with control or siRNA against ABCA1 for 48h prior to being stained with Cell Trace Far Red and overlayed onto 4T1 tumor spheroids previously stained with PKH26 (Sigma Aldrich), in an ultra-low attachment 96-well plate. Spheroids were then removed, disrupted mechanically and quantified by flow cytometry. Different letters denote a statistically significant difference ($P<0.05$, Student's T Test).

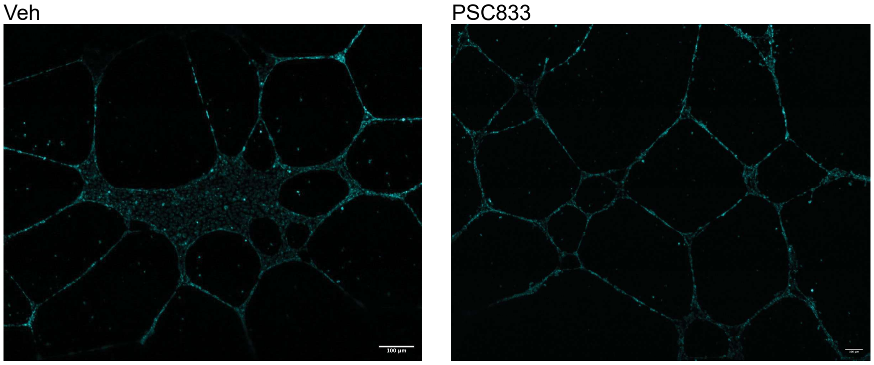


Supplementary Fig. 3

**Supplementary Figure 3:** ***ABCA1 inhibitor PSC833 modestly increases ability of macrophages to support tube formation by HUVEC cells***. Representative micrographs shown. Data is in support of **Fig. 3.**

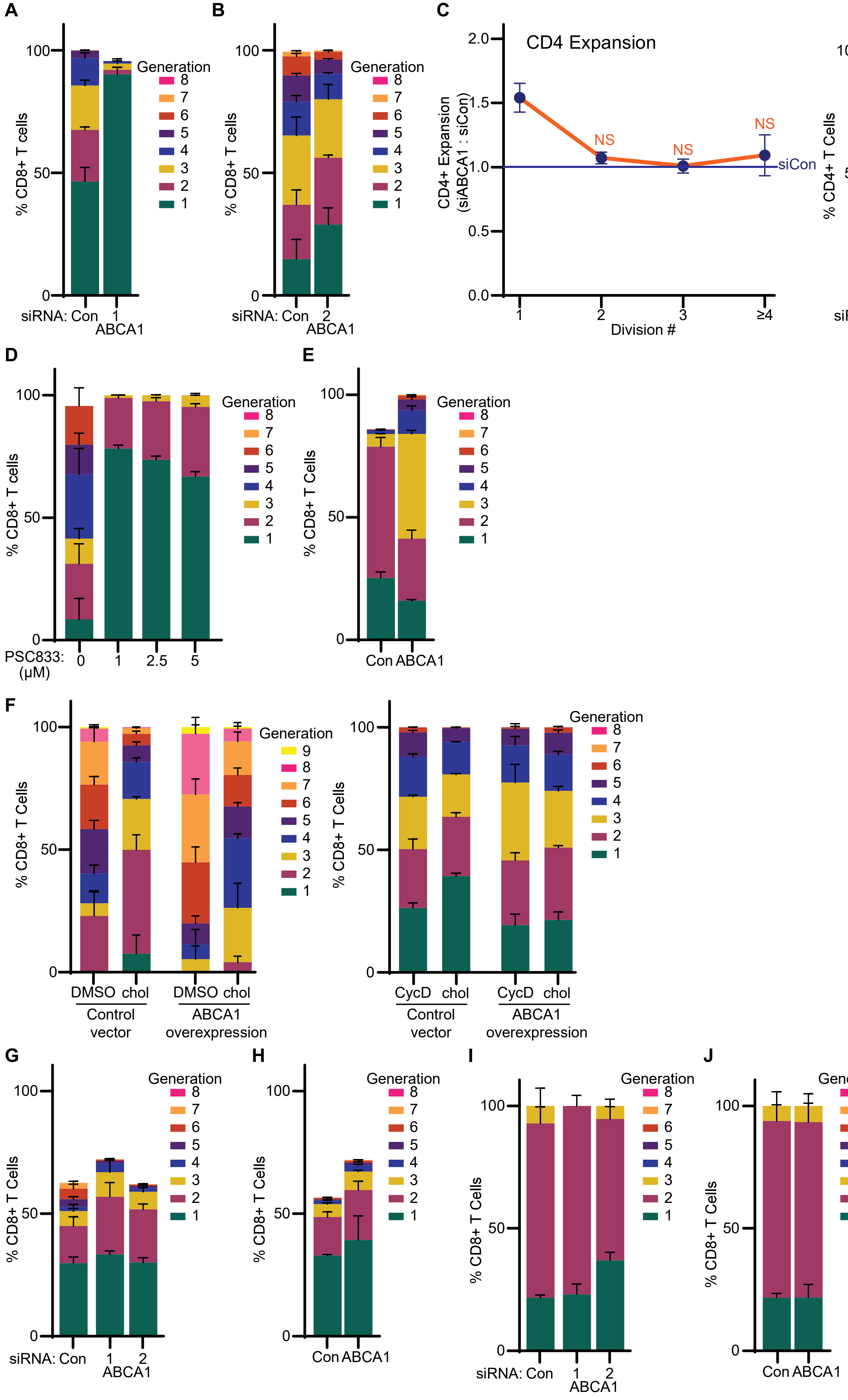


Supplementary Fig. 4

**Supplementary Figure 4:** ***ABCA1 activity results in BMDMs that support T cell migration, expansion.*** (A) CD8+ T cell expansion is decreased when activated T cells are co-cultured with BMDMs transfected with siRNA against ABCA1. The percentage in each clonal generation shown here. Data in support of **Fig. 5C**. **(B)** A second siRNA against ABCA1 yielded similar results. **(C)** The effects of siRNA against ABCA1 in BMDMs on T cell expansion were restricted to CD8+ T cells. When CD4+ T cell expansion was assessed, no significant differences were observed (same experiment as in **Fig. 5C** where CD8+ expansion is shown). A ratio between the proportion of CD4+ T cells expanded under treatment conditions to vehicle control conditions is shown to the left, with the blue line being siControl conditions. The corresponding percentage in each clonal generation is shown to the right. **(D)** CD8+ T cell expansion is decreased when activated T cells are co-cultured with BMDMs pretreated with the ABCA1 inhibitor, PSC833. The percentage in each clonal generation is shown. Data in support of **Fig. 5D**. **(E)** Overexpression of ABCA1 in BMDMs enhanced CD8+ T cell expansion. The percentage in each clonal generation shown here. Data in support of **Fig. 5E**. **(F)** BMDMs were transfected with either control or an ABCA1 overexpression vector. 48 hours post-transfection, BMDMs were incubated in 50μM cholesterol for 24 hours prior to being washed and then co-cultured with activated T cells. Left: Resulting CD8+ T cell expansion after BMDMs were treated with cholesterol solubilized in DMSO. The percentage in each clonal generation shown. Right: Resulting CD8+ T cell expansion after BMDMs were treated with cholesterol solubilized in cyclodextrin. The percentage in each clonal generation shown. **(G)** The robust decreases in CD8+ T cell expansion when pan-T cells were cultured with BMDMs lacking ABCA1 is lost when only CD8+ T cells are cultured with BMDMs. CD8+ T cells were bead-isolated prior to incubation with BMDMs that had previously been transfected with control or two different siRNAs against ABCA1. Resulting CD8+ T cell expansion was assessed. The percentage in each clonal generation shown. **(H)** Similarly, the increased CD8+ T cell expansion observed when pan-T cells are co-cultured with BMDMs overexpressing ABCA1 is muted when only CD8+ T cells are cultured with BMDMs. The percentage in each clonal generation is shown. **(I-J)** The robust effects of BMDM-ABCA1 on CD8+ T cell expansion are attenuated when contact between BMDMs and T cells is prevented through the use of a Boyden chamber. The percentage in each clonal generation after BMDMs were transfected with two different siRNAs against ABCA1 is shown in **I**. The percentage in each clonal generation after BMDMs were transfected with an ABCA1 expression vector is shown in **J**.

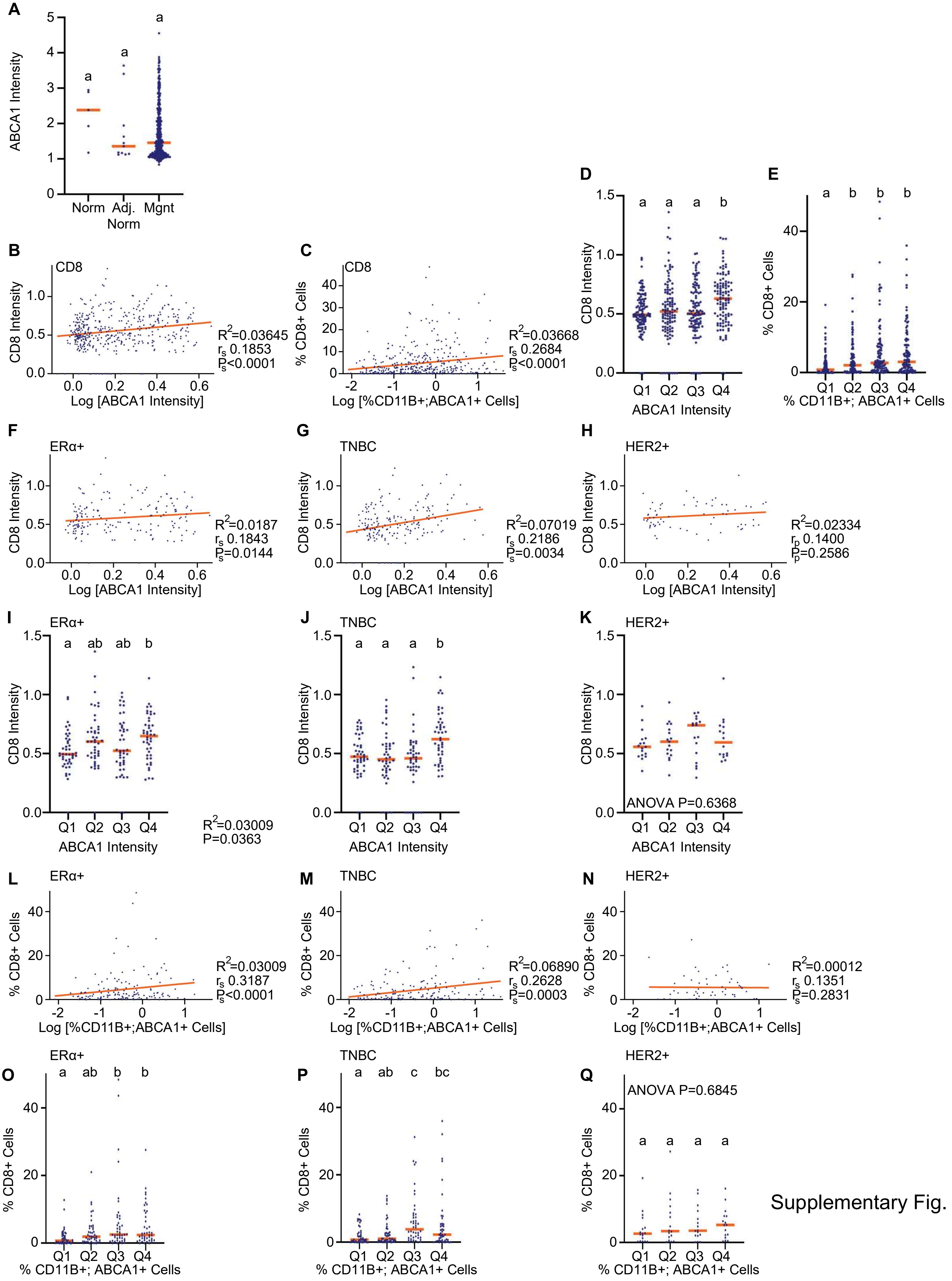


Supplementary Fig. 5

**Supplementary Figure 5:** ***ABCA1 staining is associated with CD8 staining in human breast tumor tissue microarrays.*** The data in this supplementary figure are in support of **Fig. 6**. Human breast tumor tissue microarrays (TMAs) were co-stained for ABCA1, the pan-myeloid cell marker CD11B, the cytotoxic T cell marker CD8, and the nuclear stain DAPI, in a mIF assay. Staining intensity as well as % cells positive for each stain was quantified. **(A)** A few cases of normal breast and tumor-adjacent normal breast tissue were included in some of the TMAs. Here, ABCA1 staining intensity is shown when parsed this way. We had too few samples to power the statistical analysis for this. **(B)** There is a positive correlation between ABCA1 and CD8 staining intensity (Spearman coefficient and P values shown). **(C)** There is a positive correlation between percentage of cells co-staining for CD11B and ABCA1, and the percentage of CD8+ cells. Spearman coefficient and P values shown. **(D-E**) Data for B & C were parsed based on ABCA1 into quartiles. Different letters denote statistical difference from one-another ($P<0.05$; Kruskal-Wallace followed by Dunn's multiple comparison test). **(F-H)** Samples for each different breast cancer subtypes were assessed independently, evaluating staining intensity. Spearman coefficients and P values shown for F&G while the Pearson coefficient and P value is shown for H. **(I-K**) Data for F-H were parsed based on ABCA1 into quartiles. Different letters denote statistical difference from one-another ($P<0.05$; Kruskal-Wallace followed by Dunn's multiple comparison test, with the exception of K, where a 1-Way ANOVA was used with the P value indicated). **(L-N**) Samples for each different breast cancer subtypes were assessed independently, evaluating the percentage of positive cells. Spearman coefficient and P values shown. **(O-Q)** Data for L-N were parsed based on percentage of CD11B+;ABCA1+ cells into quartiles. Different letters denote statistical difference from one-another ($P<0.05$; Kruskal-Wallace followed by Dunn's multiple comparison test, with the exception of Q, where a 1-Way ANOVA was used with the P value indicated).

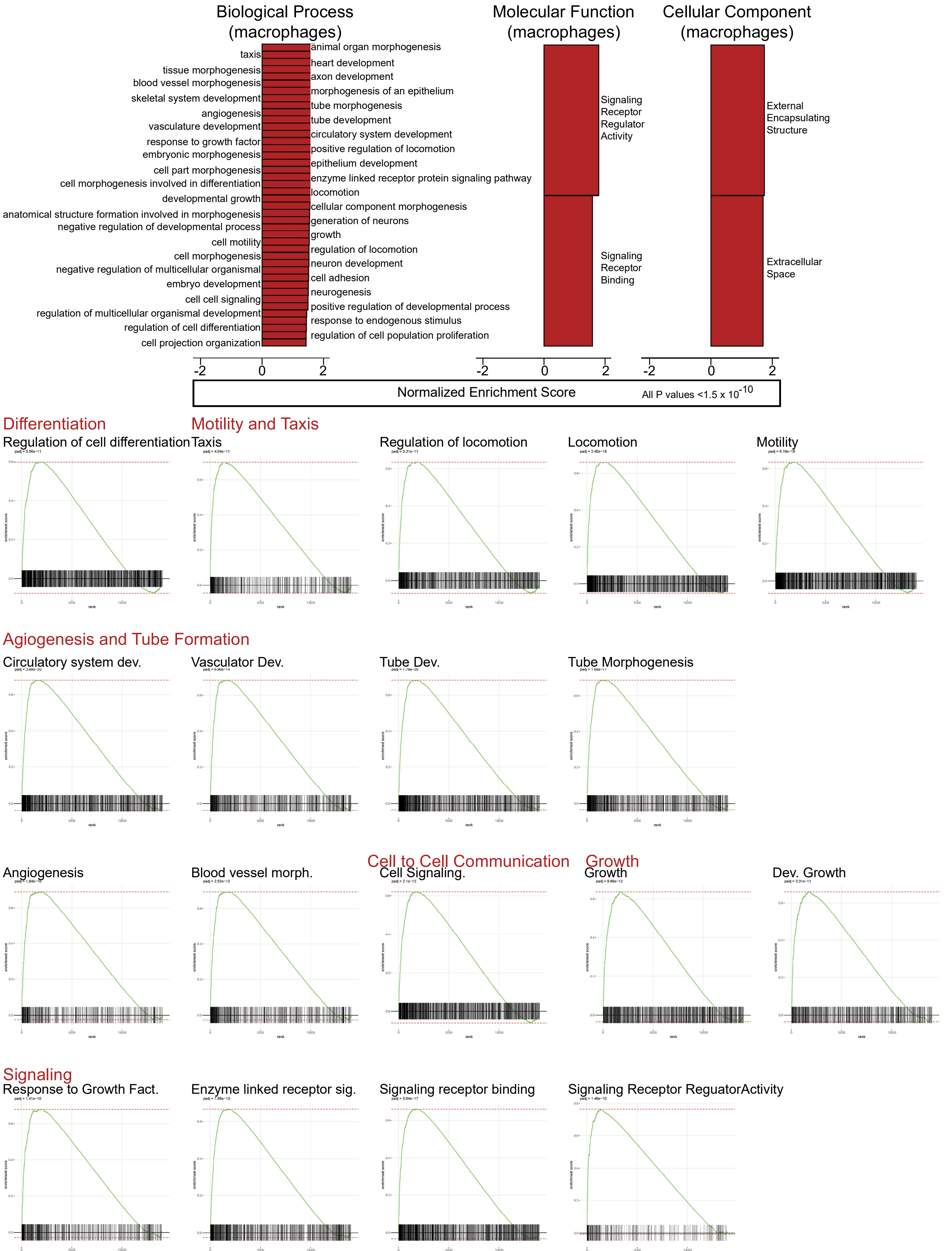


Supplementary Fig. 6

**Supplementary Figure 6:** ***GO analysis of DEGs between control and siRNA-ABCA1 transfected BMDMs***. Gene Ontology (GO) was performed on the differentially expressed genes (DEGs) identified from RNA-Seq. GO enrichment was calculated and visualized to highlight the most significantly enriched GO Terms using ShinyGO.

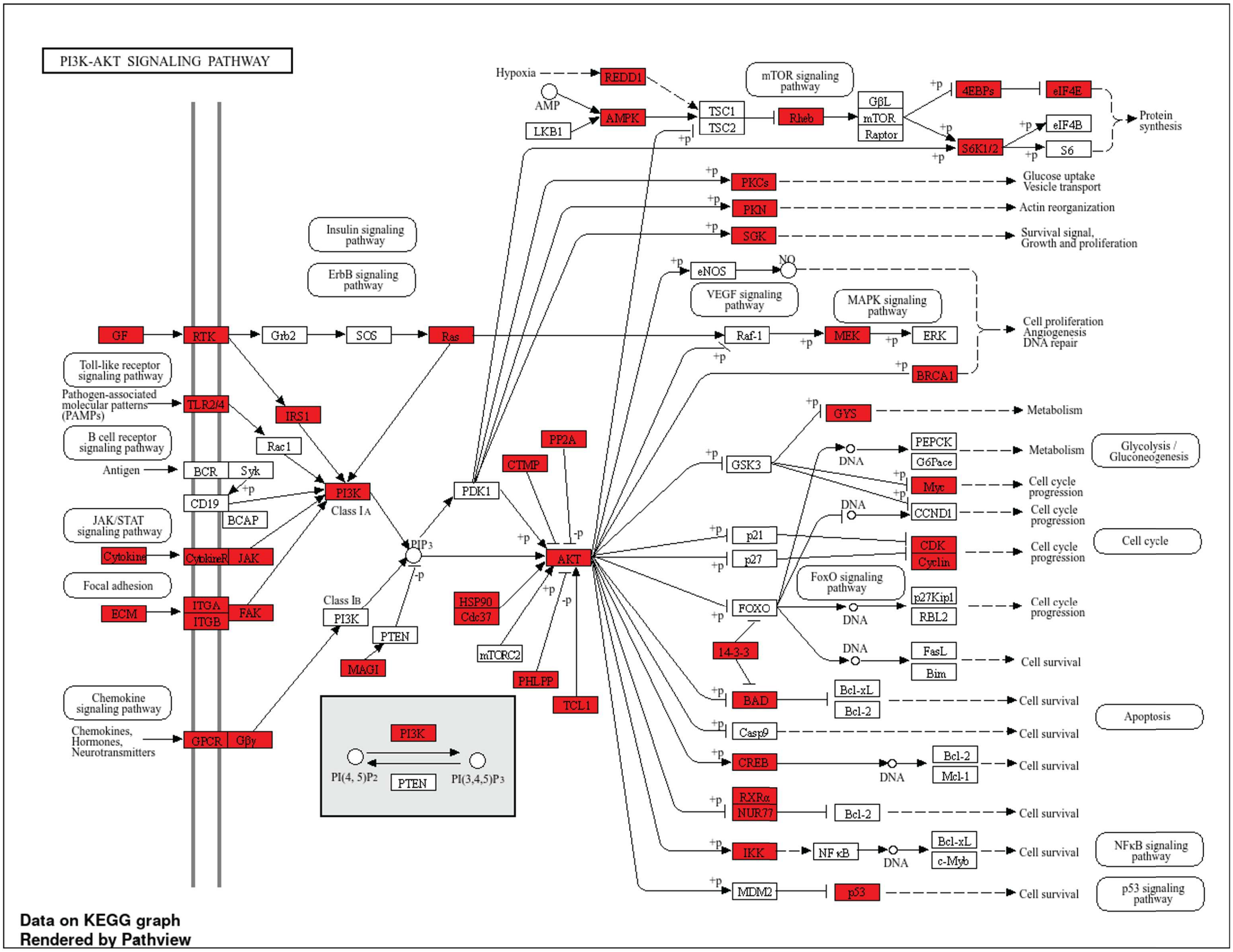


Supplementary Fig. 7

**Supplementary Figure 7:** ***KEGG Pathway analysis of DEGs DEGs between control and siRNA-ABCA1 transfected BMDMs***. Gene Set Enrichment Analysis (GSEA) for KEGG Pathway analysis was conducted on DEGs from RNA-Seq using the clusterProfiler package in R. Results were visualized using GSEA plots and enrichment maps to highlight significantly enriched KEGG pathways.

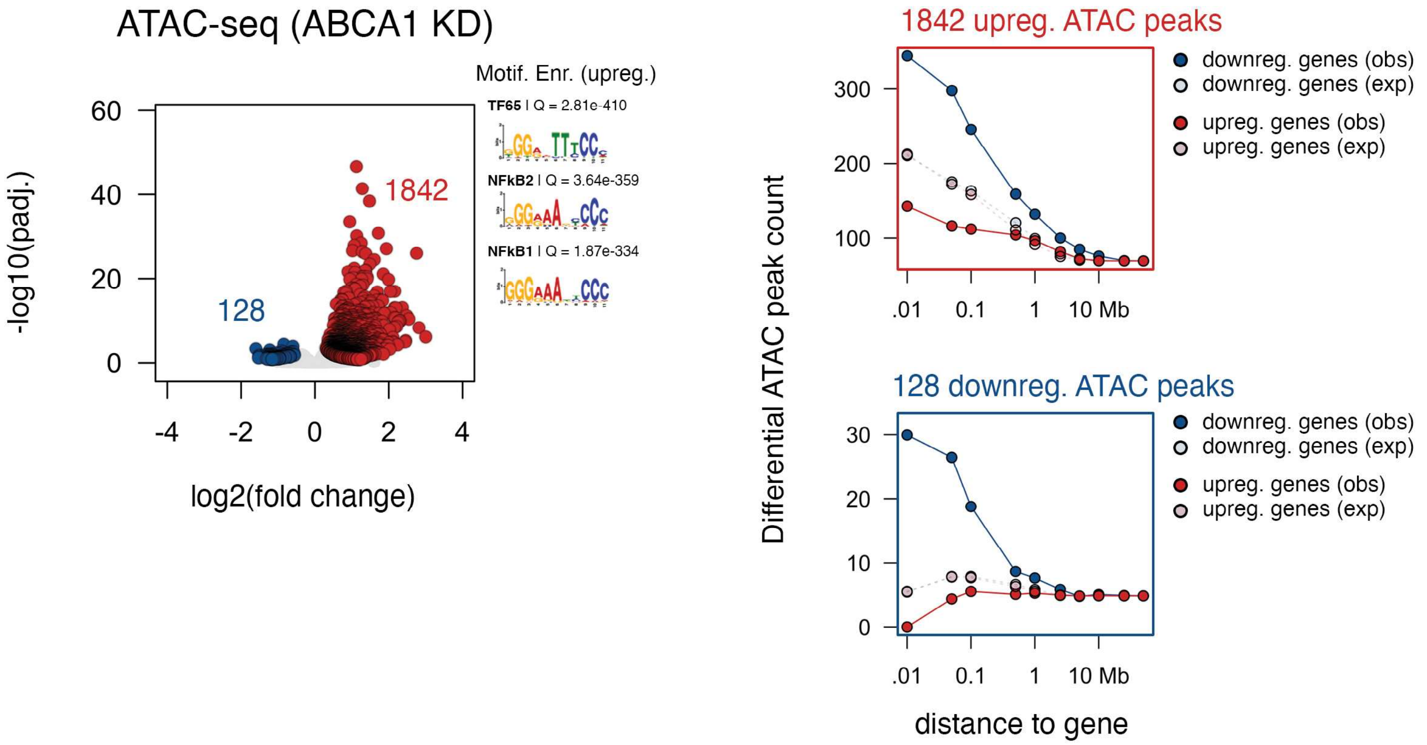


Supplementary Fig. 8

**Supplementary Figure 8: *ATAC-seq analysis of BMDMs transfected with control or siRNA against ABCA1*.** Differential ATAC-seq and enrichment proximal to dynamic gene signatures following ABCA1 depletion. Left: Volcano plot analysis of significantly upregulated (red, 1,842 peaks) and downregulated (blue, 128 peaks) following ABCA1 knockdown. Inset highlights top 3 motifs enriched in upregulated ATAC peaks (SEA analysis; MEME suite). Right: Proximity-based enrichment of upregulated ATAC peaks (top, red) and downregulated peaks (bottom, blue) relative to significant upregulated genes (red) and downregulated genes (blue). Observed densities (solid) are shown in relation to expected densities (dashed).

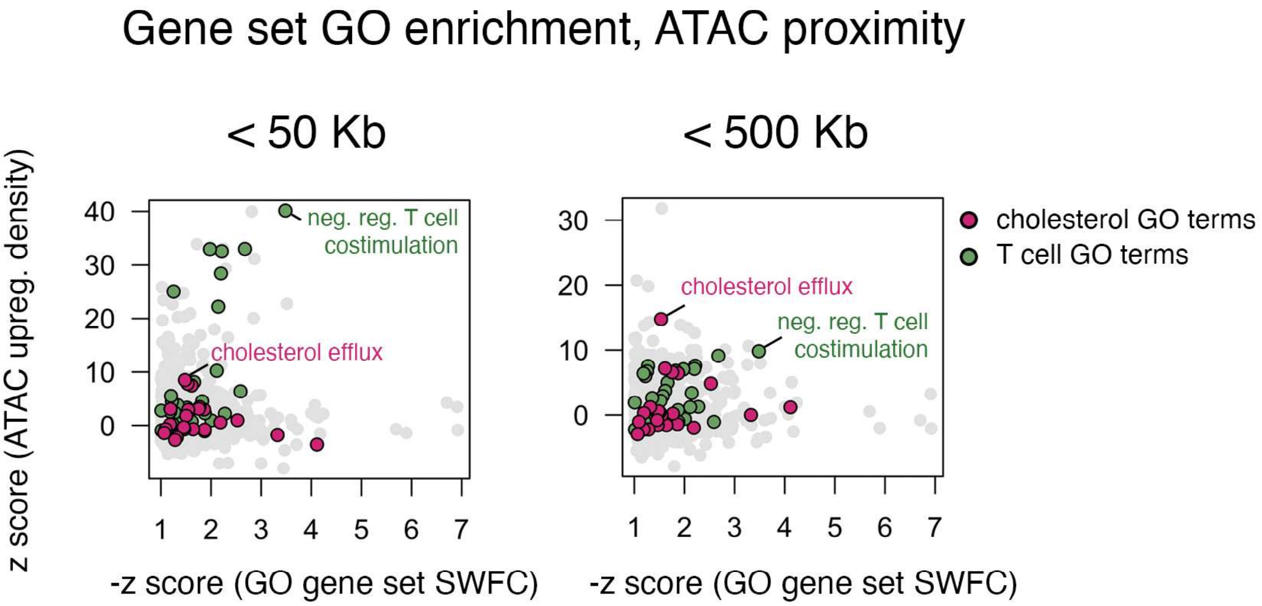


Supplementary Fig. 9

**Supplementary Figure 9:** ***ATAC-seq analysis of BMDMs transfected with control or siRNA against ABCA1***. Proximity-based enrichment of upregulated ATAC peaks relative to specific gene sets, defined by shared gene ontology (GO). Z-score ATAC density (y-axis) represents the proximal density of upregulated ATAC peaks (50 Kb, left; 500 Kb, right) compared against a randomized null model. -Z score GO gene set (x-axis) represents the mean significance weighted fold change (SWFC) for a given gene set (with shared GO term) compared against a randomized null model. GO term:: Gene sets related to cholesterol (magenta) and T cell (green) are highlighted, with top GO term enrichments for "Negative Regulation of T cell costimulation" (50 Kb) and "Cholesterol Efflux" (500 Kb) labeled.

| PWM ID | PWM Logo | Cell Type | TFs with Matching Motifs | | | | |
|---|---|---|---|---|---|---|---|
| 97 | 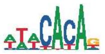 | | | | | | |
| 93 | 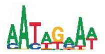 | M | MEF2C | MEF2A | MEF2D | PROP1 | PO5F1 |
| 83 | 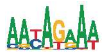 | M,T | GCR | PRGR | MEF2C | STAT1 | NFAC1 |
| 30 | 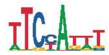 | M | STAT1 | MEF2C | MEF2A | PO2F1 | FOXL2 |
| 100 | 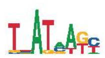 | M,T | GATA4 | GATA3 | GATA2 | | |
| 16 | 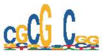 | | | | | | |
| 35 | 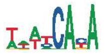 | M,T | GFI1B | GFI1 | RUNX1 | BRAC | SP7 |
| 71 | 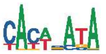 | | | | | | |
| 1 | 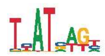 | M,T | GATA4 | GATA3 | GATA2 | GATA1 | TAL1 |
| 27 | 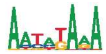 | M | FOXL2 | FOXJ3 | MEF2C | CDX1 | IRF9 |
| 32 | 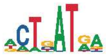 | M | GATA4 | GATA3 | GATA2 | TAL1 | |
| 79 | 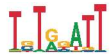 | M,T | RUNX1 | RUNX2 | RUNX3 | TWST1 | |
| 21 | 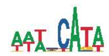 | M,T | FOXJ3 | HXA10 | SRY | TWST1 | HXA9 |
| 91 | 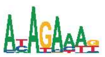 | | | | | | |
| 111 | 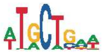 | | | | | | |
| 23 | 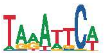 | | | | | | |
| 88 | 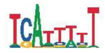 | M,T | NKX61 | STAT2 | NFAC1 | MAFF | ONEC2 |
| 109 | 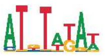 | | | | | | |

Supplementary Fig. 10

**Supplementary Figure 10: SEAMoD *Nueral network learning using RNAseq and ATACseq datasets identified transcription factors likely to drive the changes in differentially expressed genes.*** Motifs obtained from the SEAMoD model trained on the data from the macrophages (M) and co-cultured T cells (T). Convolutional filters were converted into position weight matrices (PWMs), which were then matched with a compendium of previously characterized motifs in mouse using TOMTOM software. Only those motifs were selected that were found significant by the model in one or both cell types and had a strong match with known mouse motifs (see methods). Note that only top 5 Mouse motif matches are shown here. Missing motifs indicate that either no or fewer than 5 matching motifs were found.

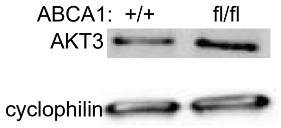


Supplementary Fig. 11

**Supplementary Figure 11:** ***AKT3 protein expression is increased in BMDMs lacking ABCA1***. Representative Western blot AKT3 comparing BMDMs from ABCA1$^{+/+}$;LysMCre$^{+}$ mice to BMDMs from ABCA1$^{fl/fl}$;LysMCre$^{+}$ mice.
